# Classification of hospital and urban wastewater resistome and microbiota over time and their relationship to the eco-exposome

**DOI:** 10.1101/697433

**Authors:** Elena Buelow, Andreu Rico, Margaux Gaschet, José Lourenço, Sean P. Kennedy, Laure Wiest, Marie-Cecile Ploy, Christophe Dagot

## Abstract

Wastewaters (WW) are important sources for the dissemination of antimicrobial resistance (AMR) into the environment. Hospital WW (HWW) contain higher loads of micro-pollutants and AMR markers than urban WW (UWW). Little is known about the long-term dynamics of H and U WW and the impact of their joined treatment on the general burden of AMR. Here, we characterized the resistome, microbiota and eco-exposome signature of 126 H and U WW samples treated separately for three years, and then mixed, over one year. Multi-variate analysis and machine learning revealed a robust signature for each WW with no significant variation over time before mixing, and once mixed, both WW closely resembled U signatures. We demonstrated a significant impact of pharmaceuticals and surfactants on the resistome and microbiota of H and U WW. Our results present considerable targets for AMR related risk assessment of WW.

---

The worldwide spread of multidrug-resistant bacteria is an important public health issue with a high health and economic burden^1–3^. A global “One Health” approach is urgently needed to combat the dissemination of antibiotic resistant bacteria (ARB) and antimicrobial resistance genes (ARGs) from humans and livestock to the environment and vice versa, as well as to identify key drivers contributing to the selection, dissemination and persistence of ARB and/or ARGs^4,5^. The natural environment and its biodiversity serve as a wide reservoir of genetic determinants implicated in resistance to antimicrobial compounds^6,7^. Human activity has a significant impact on the terrestrial and aquatic microbial ecosystems through chemical pollutants that are spread via urban, agricultural and industrial waste and which pose an important selective pressure for AMR^8^. For instance, urban and hospital wastewaters (UWW and HWW) contain a high diversity of ARGs and chemicals^9–12^. It is therefore generally accepted that the implementation of efficient wastewater treatment plants (WWTP) is essential in order to reduce the amounts of chemicals, ARGs and ARB that reach the environment^4,9,13^. The treated WW are re-introduced into the aquatic environment and the produced sludge often re-used in agricultural lands^11,14^. However, despite a global reduction of ARGs through treatment, effluents from urban, hospital and industrial wastewater still contain ARGs, antibiotics (ABs) and moderate levels of other pollutants affecting microorganisms (e.g. biocides, heavy metals)^11,15,16^. HWWs have been reported to contain particularly high amounts of ARGs, ARBs and ABs^9,12,17^. It has been debated whether HWW contributes significantly to the load of ARGs in the UWW systems, and whether separate treatment for HWWs should be applied^9,13,18^. Recent work has shown that HWW has limited impact on the relative levels of ARGs and mobile genetic elements (MGEs) associated with ARGs like integrons, in hospital receiving urban wastewater (WW)^9,15,19^. However, most studies usually analyzed a limited number of samples and yet, longitudinal studies that monitor WW dynamics are so far lacking, which limits the possibility of assessing the risk for AMR mediated through WW.

We thus studied the dynamics of the resistome, microbiota and of some chemical-pollutants present in WW (the “eco-exposome”) of 126 WW samples (UWW, HWW and mixed WW) in a French city during a period of approximately four years: 34 months with separate treatments for H and U WW and 11 months with H and U WW mixed. We studied H and U WW in the course of their passage through i) two independent wastewater treatment systems applying the conventional secondary (activated sludge) treatment process and ii) then mixed 1:2 (HWW:UWW) into one system. We investigated the relationship of the respective resistome and microbiota with the measured eco-exposome (pharmaceuticals, mainly antibiotics; surfactants and heavy metals). Multi-variate analysis and machine learning, reveal a robust signature of the resistome, microbiota and eco-exposome of HWW compared to UWW with no significant variation over time. We also showed that, when mixed, both WW closely resemble urban signatures. Furthermore, we demonstrated that pharmaceuticals and surfactants had a large influence on the variability of the monitored resistome and microbiota of H and U WW.

## Results

### HWW and UWW have distinct resistome and microbiota signatures

We evaluated the resistome and microbiota of monthly WW (N=126) and river (N= 12) samples. For the resistome, we targeted 78 genes by high-throughput qPCR^9,20^ conferring resistance to antibiotics, quaternary ammonium compounds, or heavy metals, grouped into 16 resistance gene classes. The genes targeted include ARGs that are most commonly detected in the gut microbiota of healthy individuals^21,22^, clinically relevant ARGs (including genes encoding extended spectrum β-lactamases (ESBLs), carbapenemases, and vancomycin resistance), and heavy metal and quaternary ammonium compound resistance genes suggested to favor cross and co – selection for ARGs in the environment^23,24^. We also targeted genetic elements as important transposase gene families^25^ and class 1, 2 and 3 integron integrase genes, that are important vectors for ARGs in the clinics and often used as proxy for anthropogenic pollution^26^ (gene targets and grouping of genes into gene classes/according to function are detailed in Supplementary Tables 5 and 6).

H and U WW samples were treated separately through 2012 and 2014, and were mixed at a ratio of 1:2 (HWW:UWW) throughout the year 2015 (Figure 1). H and U WW samples exhibited a distinct signature with respect to the proportional makeup of their resistome (Figure 2a) and microbiota (Figure 2b). Analyzing the data with a Random Forest machine learning approach showed that the distinct H and U WW signatures resulted in a high prediction accuracy. When using the resistome as predictor (on the level of gene classes), 93.5% and 96.7% of untreated HWW and UWW samples respectively, could be correctly classified (Supplementary Figure 1a). We further analyzed the data on the individual gene level to increase resolution of the machine learning approach. Using individual genes resulted in similarly high predictions (93.5% prediction for untreated HWW and 100% prediction for untreated UWW) (Supplementary Figure 1b). Similarly, when using the microbiota, 96.8% of untreated HWW and 89% of untreated UWW samples were correctly classified (Supplementary Figure 1c).

**Figure 1:**
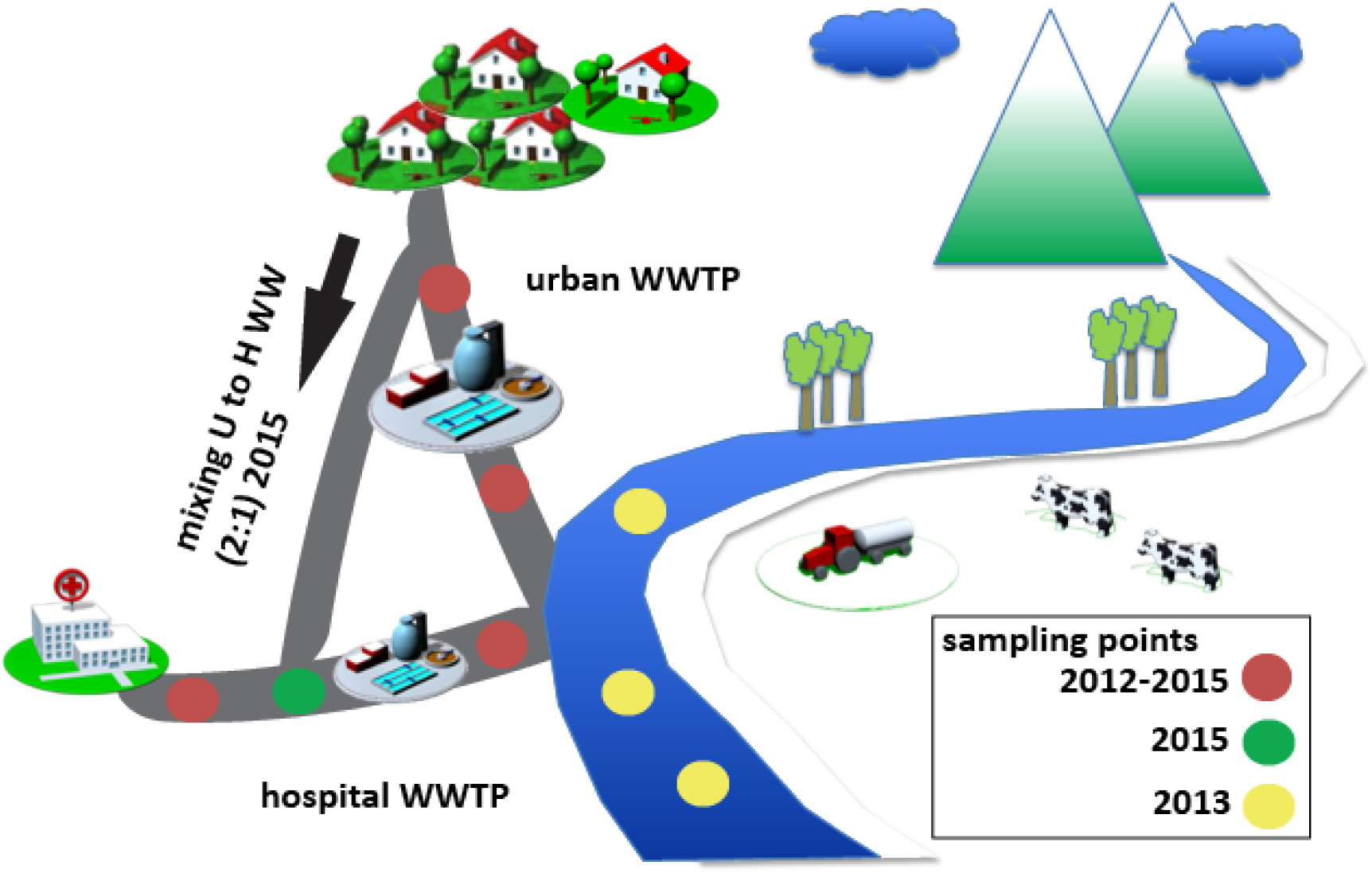
Sampling site. Samples were collected in monthly intervals (untreated and treated samples) by flow proportional sampling, from March 2012 through November 2015. From March 2012 to December 2014, UWW and HWW were treated by separate wastewater treatment plants (WWTPs). During the period from January 2015 through November 2015, UWW was mixed into the HWW (1:2 ratio HWW:UWW) and added to the separate HWW treatment line resulting in mixed WW (MWW). In addition, 12 water samples of the effluent receiving river up (river upstream) and downstream (river downstream sampling point 1 and 2) of the effluent release pipes were collected during the winter months of 2013 (January, February, November, and December).

**Figure 2:**
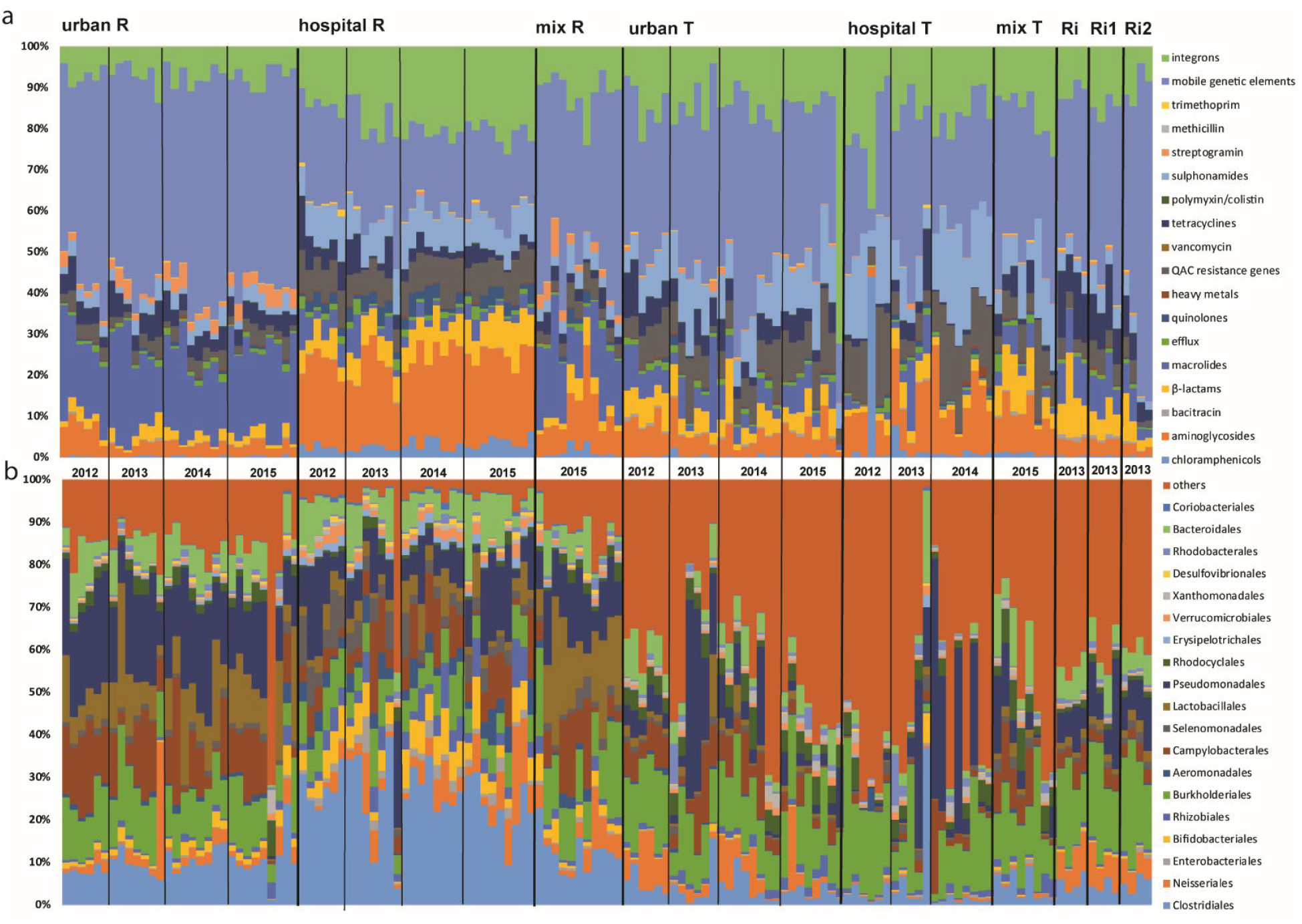
Proportional abundance of the resistome and microbiota in untreated (R) and treated (T) HWW, UWW and MWW, as well as river samples up (Ri) and downstream (Ri1 and Ri2) of the waste water release pipes throughout the sampling period (2012-2015). **a:** proportional abundance of resistome (ARG classes, heavy metals integron integrase genes and mobile genetic elements) for all samples. **b**: proportional abundance of the microbiota (displaying the 20 most abundant bacteria at the order level for all samples, where “others” represents the percentage of the remaining taxa).

For the treated H and U WW the machine learning prediction accuracy was lower compared to the untreated WW sources but still considerably high for all predictor levels and in particular for the microbiota (≥80%) (Supplementary Figure 2).

For the MWW overall classification success was lower (Supplementary Figure 1 and 2), given a much lower sample size and likely due to the mixing of the two wastewater sources hampering clear signatures.

### Temporal dynamics of H and U WW

Redundancy analysis was performed to assess putative significant influences of time on the variability of resistome and microbiota compositions using sampling year or season as independent variables (Monte Carlo (MC) permutations (n=499)). Analysis was carried out for all different sample groups together (untreated U, H, M WW and treated U, H, M WW) to potentially identify general patterns influencing all groups at the same time. The same analysis was also performed per individual sample group (untreated and treated H, U and M WW alone, respectively).

For untreated HWW and UWW, neither year (*p*=0.6 resistome and *p*=0.4 microbiota respectively) nor season (*p*=0.9 and *p*=0.3) had a significant impact on the resistome (resistance gene classes) and microbiota composition of the WW over the first 3 years before mixing (Supplementary Table 1a). However, by analyzing the sample groups separately, a yearly and seasonal impact on the level of individual resistance genes in HWW was demonstrated (*p*=0.004 year, *p*=0.032 season; Supplementary Table 1b). Redundancy analysis revealed the relationship between individual genes and gene classes with seasons, pointing towards a correlation of increased normalized abundance of individual genes and gene classes detected during summer season (Supplementary Figure 3a, Supplementary Figure 4). The fact that the hospital was only installed in February 2012^27^ may explain why the year 2012 is particular for HWW resistome, with overall lower normalized abundance of the resistome and no obvious trend towards the summer season in 2012. After HWW and UWW mixing (in 2015), no significant variation for the resistome and microbiota composition throughout the seasons could be observed considering all sample groups (untreated MWW, HWW and UWW; *p*=0.6 resistome and *p*=0.07 microbiota respectively).

For treated WW, the analysis exhibited more variation of the resistome and microbiota composition compared to the untreated sources over the years (2012-2014 for H and U WW; Figure 2, Supplementary Table 1c). Redundancy analysis revealed that in particular the microbiota of treated H and U WW effluents varied between the years (*p*=0.016 for all groups, *p*=0.021 for treated HWW and *p*=0.006 for treated UWW, Supplementary Table 1d) while no significant seasonal impact could be observed. The resistome varied between the years for treated HWW on both the gene class and the individual gene levels (*p*=0.004 and *p*=0.012, Supplementary Table 1d), whereas for the treated UWW this variation was only significant on the individual gene level (*p*=0.028). For the treated mixed WW (MWW), significant seasonal variation was observed for the microbiota composition (*p*=0.02), while for the resistome no significant variation could be observed.

### The untreated HWW resistome is significantly diluted by UWW in mixed WW

The bacterial biomass (absolute copy numbers of 16S rRNA genes per liter of water) was comparable for the untreated WW sources (HWW, UWW and MWW) (Supplementary Figure 5). Exemplary, here ratios of HWW over UWW and HWW over MWW were calculated based on the averaged cumulative abundance of the resistome in untreated HWW and UWW (Table 1).

**Table 1:** Average fold changes for gene classes cumulative abundance of untreated HWW over UWW (2012-2015; significant differences indicated by asterisk *; *p* values ≤ 0.004) and HWW over MWW (2015, significant differences indicated by asterisk *; *p* values ≤ 0.03) ± Standard Deviation. Significant differences were calculated by comparing the normalized cumulative abundance values of individual gene classes for all samples belonging to each sample group using the non-parametric Mann-Whitney test. Fold changes were calculated for individually paired samples for each gene class / sample group. NA indicates that gene classes were undetectable in either one or both of the sample groups.

| Gene classes conferring resistance to: | Fold change untreated Hospital WW/Urban WW | Fold change untreated Hospital WW/Mixed WW |
| --- | --- | --- |
| *chloramphenicol* | 84 ($\pm 93$)* | 13 ($\pm 11$)* |
| *aminoglycosides* | 43 ($\pm 31$)* | 8 ($\pm 6$)* |
| *bacitracin* | 8 ($\pm 13$)* | 7 ($\pm 4$)* |
| *beta-lactams* | 26 ($\pm 22$)* | 9 ($\pm 6$)* |
| *macrolides* | 1 ($\pm 1$) | 0.6 ($\pm 0.3$) |
| *(multi-drug) Efflux* | 9 ($\pm 11$)* | 4 ($\pm 4$)* |
| *quinolones (qnr)* | 161 ($\pm 326$)* | 10 ($\pm 8$)* |
| *heavy metals* | 7 ($\pm 9$)* | 4 ($\pm 3$)* |
| *quaternary ammonium compounds QACs)* | 18 ($\pm 14$)* | 7 ($\pm 4$)* |
| *vancomycin* | 12 ($\pm 35$)* | 18 ($\pm 20$)* |
| *tetracycline* | 4 ($\pm 3$)* | 3 ($\pm 2$)* |
| *polymixin* | 8 ($\pm 9$)* | 4 ($\pm 4$)* |
| *sulphonamides* | 19 ($\pm 12$)* | 7 ($\pm 4$)* |
| *methicillin* | NA (undetectable in UWW) | NA (undetectable in all but one MWW sample) |
| *streptogramin* | 0.2 ($\pm 0.2$)* | 0.2 ($\pm 0.2$)* |
| *trimethoprim* | 8 ($\pm 6$)* | 5 ($\pm 3$)* |
| **Gene classes grouped according to function:** |  |  |
| *transposase genes (MGEs)* | 3 ($\pm 1$)* | 2 ($\pm 1$) |
| *integron integrase genes* | 16 ($\pm 9$)* | 6 ($\pm 3$)* |

The untreated HWW contained significantly more gene classes compared to the untreated UWW, between 3 (transposase genes) and 161-fold (*qnr* genes encoding quinolone resistance) higher (*p* values ≤ 0.004). When WW were mixed at the experimental ratio of 1:2 (HWW:UWW), the untreated MWW contained significantly less resistance gene classes compared to untreated HWW, between 3 and 22-fold lower (*p* values ≤ 0.03). Interestingly, there was no significant difference for the genes encoding resistance to macrolides for both HWW over UWW and HWW over MWW comparisons (Table 1, Figure 3a and 3b). The only resistance gene significantly lower in HWW compared to UWW or MWW (*p*<0.0001), was the streptogramin resistance gene *vatB*. The *mecA* gene encoding resistance to methicillin was undetectable in all UWW and in all but one MWW samples.

**Figure 3:**
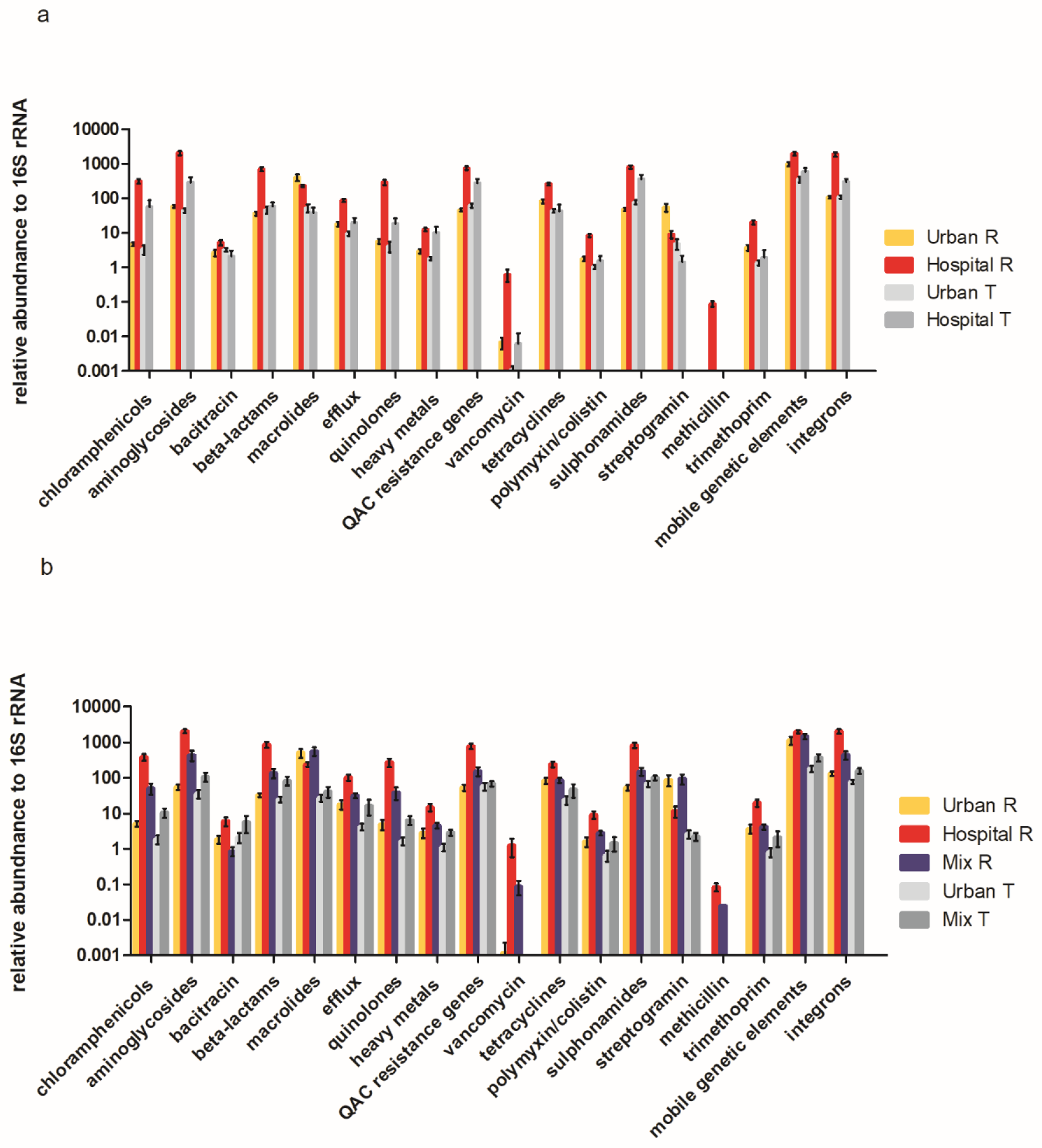
Averaged normalized abundance of ARG classes, heavy metals, MGEs and integrons over all collected samples per sample type +-standard deviation. **a**: relative abundance of ARG classes, heavy metals, QACs, MGEs and integrase genes in untreated (n21) and treated HWW (n=19), untreated (n=21) and treated (n=20) UWW averaged over the numbers of samples collected for each water type in the given time interval (March 2012 – December 2014) +-standard deviation. **b**: averaged normalized abundance of ARG classes, heavy metals, QACs, MGEs and integrase genes in untreated (n=10) HWW, untreated (n=9) and treated (n=8) UWW and untreated (n=10) and treated (n=8) MWW (at the experimental ratio of 1/3 HWW to 2/2 UWW) WW samples averaged over the numbers of samples collected for each sample group in the given time interval (January 2015-November 2015) +-standard deviation.

Altogether, these data indicate a significant dilution impact of UWW on the normalized abundance of the targeted resistome of HWW when mixing at the experimental ratio of 1:2 (HWW:UWW).

### Resistome reduction through WW treatment

The bacterial biomass (copies of 16S rRNA / liter) for all WW sources was decreased by 2-3 log after WW treatment (Supplementary Figure 5). To estimate the impact of WW treatment on the resistome, fold changes of untreated over treated H, U and M WW were calculated. The normalized cumulative abundance of all gene classes significantly decreased and was between 78 times (for genes conferring resistance to quinolones) and 5 times (for genes conferring resistance to QACs, sulphonamides and genes encoding transposase genes) lower in the treated HWW compared to untreated HWW (*p*<0.003) (Table 2; Figure 3a and 3b). When comparing untreated UWW to treated UWW, we showed a significant reduction (*p*<0.05) in the normalized cumulative abundance for 9 resistance gene classes with fold changes between 43 (for the streptogramin resistance gene *vatB*) and 3 (for genes encoding resistance to aminoglycosides) times (Table 2, Figure 3a and 3b). No significant reduction was observed after treatment of UWW for gene classes conferring resistance to bacitracin, beta-lactams, quinolones, heavy metals and quaternary ammonium compounds, and for genes encoding integron integrases. Surprisingly, sulphonamide resistance encoding genes were found to be significantly enriched after UWW treatment (*p*<0.05) (Table 2, Figure 3a and 3b). For MWW, a similar removal efficacy as for UWW could be observed (Table 2, Fig. 3b), with significant decrease for the normalized cumulative abundance of 10 genes classes and with fold changes between 41 (for the streptogramin resistance gene *vatB*) and 3 (for the genes encoding integron integrase genes) times. No significant decrease for gene classes conferring resistance to bacitracin, beta-lactams, tetracycline, heavy metals, QACs and sulphonamides could be detected.

**Table 2:**
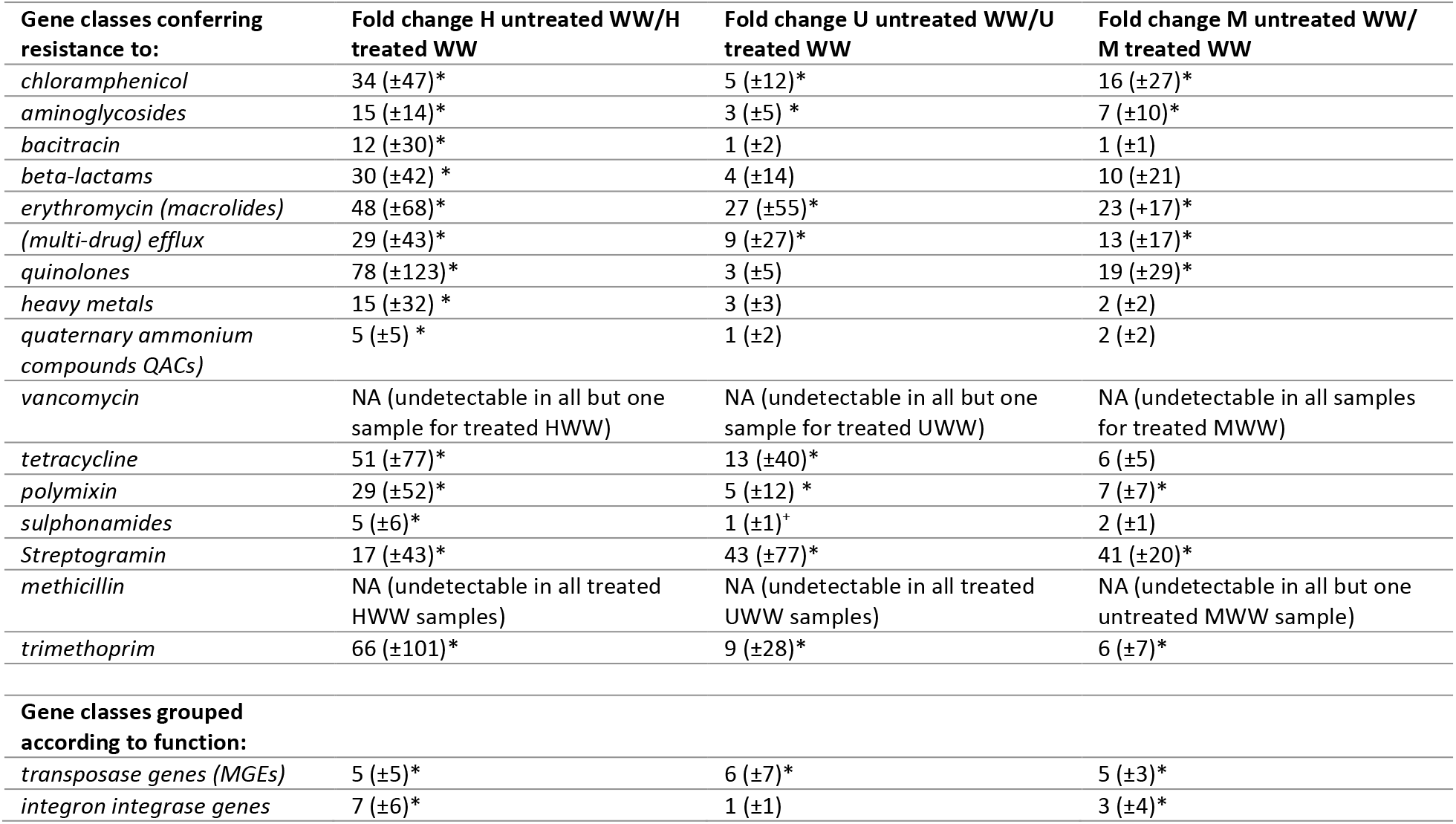
Average fold changes for gene classes of untreated WW over treated WW for H, U and M WW (*p* values ≤ 0.03) ± Standard Deviation. *= significantly lower; + = significantly higher. Significant differences were calculated by comparing the normalized cumulative abundance values of individual gene classes for all samples belonging to each sample group using the non-parametric Mann-Whitney test. Fold changes were calculated for individually paired samples for each gene class / sample group. Average fold change ± Standard Deviation are depicted in the table for comparison. NA indicates that gene classes were undetectable in either one or both of the sample groups.

### No evident impact of treated hospital WW on the receiving river

We also analyzed 12 river samples up and downstream the WWTP to evaluate putatively associated risks with the release of the treated WW into the effluent receiving river and downstream environment (Supplementary Figure 6). The resistome for the river samples collected for the sites up (Ri) and downstream (Ri1 and Ri2) of the effluent release pipes during winter season 2013 was not significantly different for either of the three sampling sites (Supplementary Figure 6). There was no significant difference of the relative abundance for any of the detected gene classes in the river samples compared to treated UWW. On the contrary, the relative abundance of nine resistance gene classes including genes encoding MGEs and integron integrase genes was significantly lower (*p*<0.04) in river samples when compared to treated HWW (Supplementary Figure 6).

### Human gut bacteria are enriched in HWW

The human gut microbiota is an important reservoir of ARGs^20,28,29^ and bacteria of the human gut are likely to be shed into the environment via wastewaters that contain at least partially human feces. We calculated the relative abundance of anaerobic human gut bacteria, as well as Enterobacteriales in the respective WW. Enterobacteriales were specifically detected as many of these Gram-negative bacteria are also pathogens. The orders Clostridiales, Bifidobacteriales and Bacteroidales which represent the most important and abundant anaerobic human gut bacteria^30,31^, were grouped together and are referred to as anaerobic human gut bacteria. Untreated HWW contained significantly higher levels of anaerobic human gut bacteria (38% ± 11 standard deviation) and Enterobacteriales (2% ± 1.5) compared to all other samples (Figures 4a and 4b). Interestingly, these orders are comparable in their relative abundance for untreated UWW and MWW, indicating a significant dilution effect of UWW in HWW (Figure 4b), as observed for the resistome. The treatment allowed a significant (p<0.05) decrease of the relative abundance of these orders for HWW and UWW (Figure 4a and 4b) whereas the reduction for MWW was only marginally significant (p=0.05) (Figure 4b).

**Figure 4:**
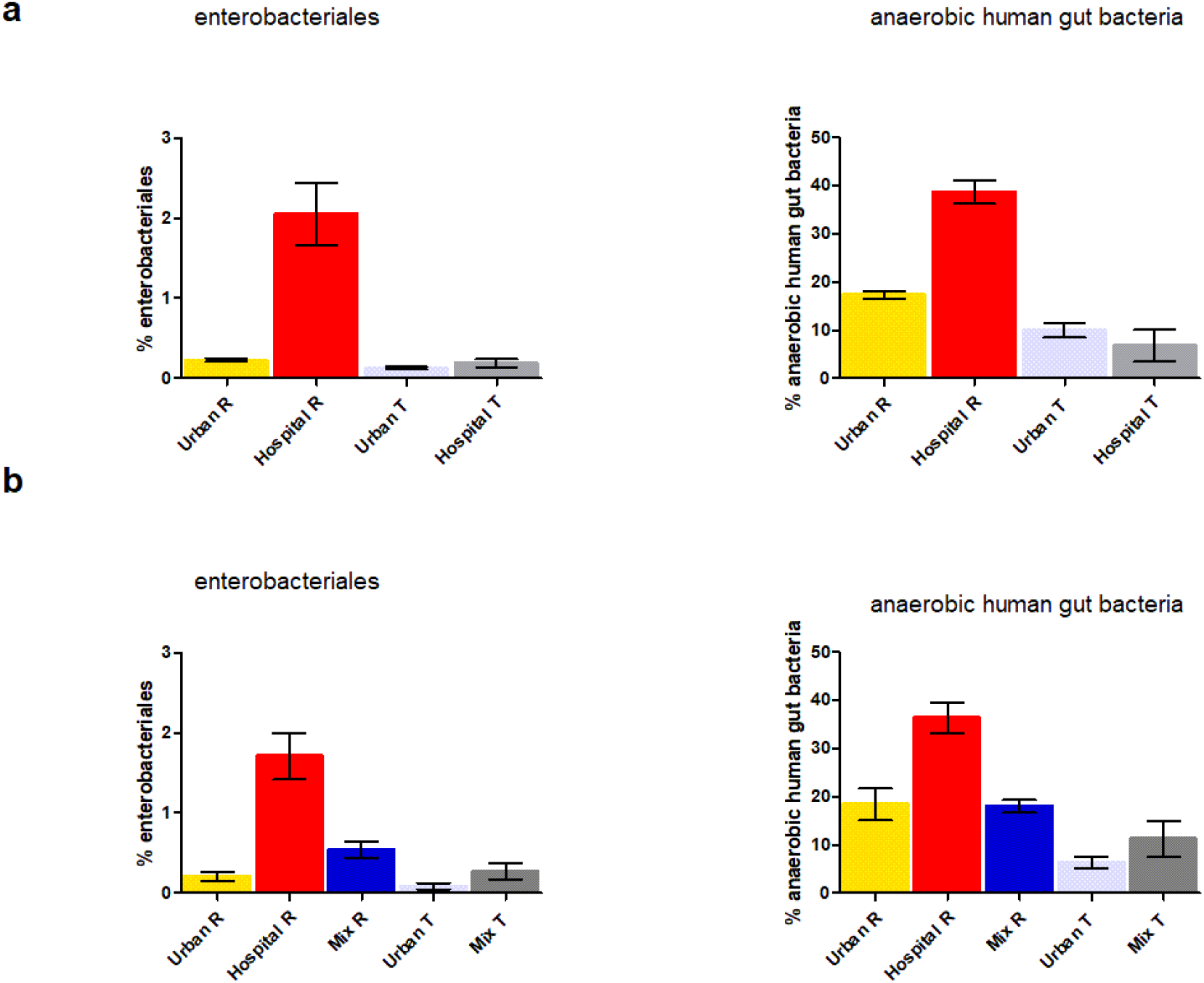
**a: Relative abundance of anaerobic human gut bacteria (Clostridiales, Bifidobacteriales and Bacteroidales) and Enterobactertiales** in untreated (n=21) and treated HWW (n=19), UWW (n=21) and (n=20) averaged over the numbers of samples collected for each sample group between March 2012 and December 2014 +-standard deviation. **b**: relative abundance of anaerobic human gut bacteria and Enterobactertiales in untreated (n=10) HWW, untreated (n=9) and treated (n=8) UWW and untreated (n=10) and treated (n=8) MWW samples (at the experimental ratio of 1:2 HWW:UWW) averaged over the numbers of samples collected for each sample group between January 2015 and November 2015 +-standard deviation.

### The eco-exposome plays an important role in shaping the resistome and microbiota in hospital and urban WW

In order to estimate the eco-exposome in the WW, selected pharmaceutical and chemical compounds including antibiotics, surfactants and heavy metals were quantified in untreated WW (Supplementary Table 2).

The relationship between the resistome/microbiota and the eco-exposome was visualized by means of PCA biplots. Here, HWW and UWW form two distinct clusters, while the MWW clusters closely to the UWW for both resistome and microbiota (Figure 5a and 5b). Antibiotics, non-ionic and cationic surfactants are the most important contributors to the respective HWW resistome and microbiota (length of arrows), while anionic surfactants and some metals are more associated with the resistome and microbiota of the UWW (Fig. 5a and 5b). Moreover, the putative impact of the measured chemicals (eco-exposome) on the resistome and microbiota variation was statistically assessed by RDA. The results of the Monte-Carlo permutations indicated that the eco-exposome significantly influences the resistome and microbiota (p-values 0.002).

**Figure 5:**
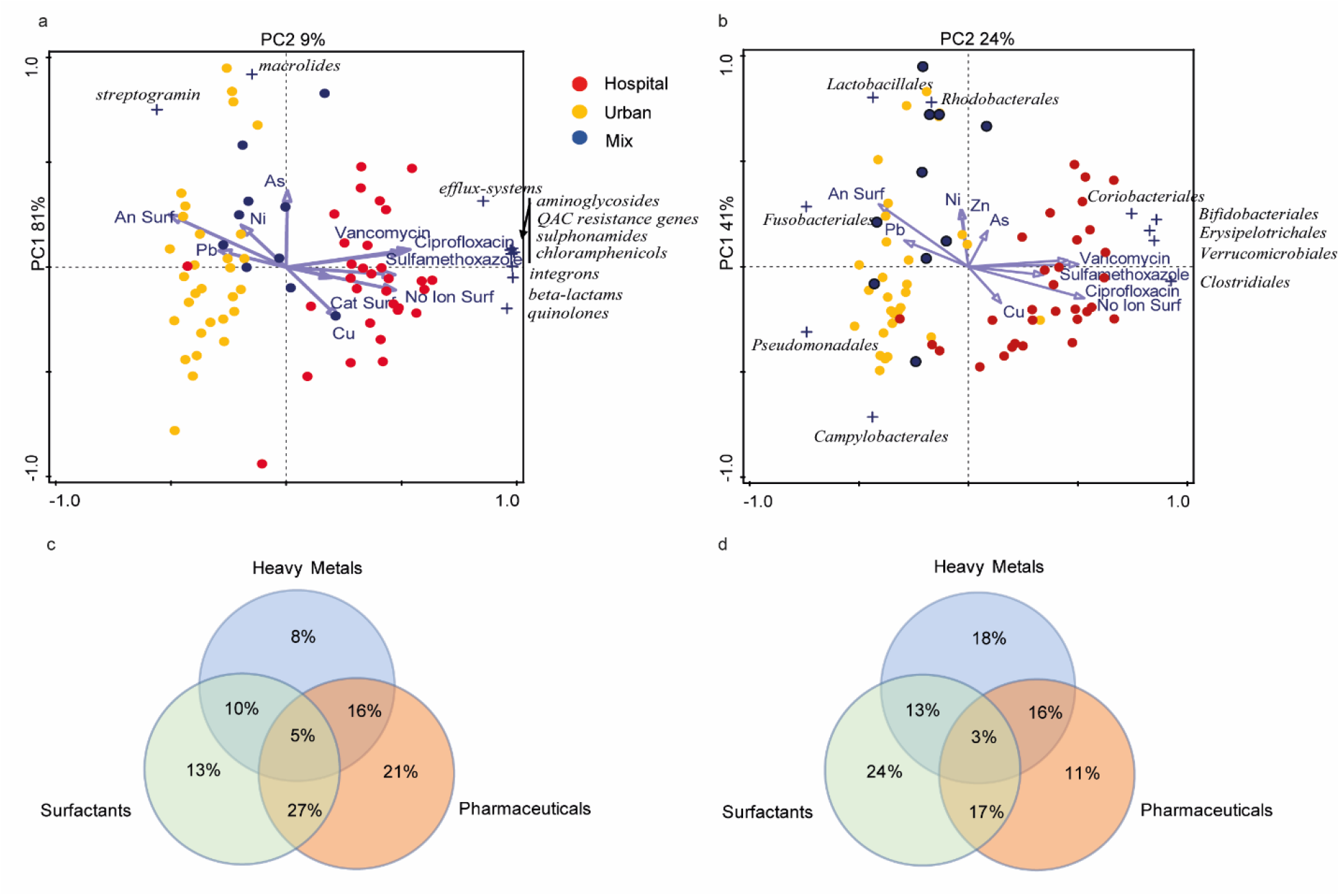
Principal component analysis showing the relationship between the eco-exposome (heavy metals, pharmaceuticals and surfactants) and the resistome (a) and microbiota (b) of untreated HWW, UWW and MWW samples; and Venn diagrams showing the results of the variation partitioning analysis with the different measured chemical classes and the resistome (c) and microbiota (d). In the PCA analysis, dots refer to urban (yellow), hospital (red) and mixed (blue) untreated WW samples.

Finally, a variation partitioning analysis was performed to study which group of the measured compounds (heavy-metals, pharmaceuticals, surfactants) might have a larger contribution on the resistome and microbiota variation, and to explore whether the interaction between these compounds has a stronger influence than the individually grouped compounds (Figure 5c and 5d). Pharmaceuticals (the antibiotics ciprofloxacin, sulfamethoxazole and vancomycin, and the neurological drug carbamazepine) explain the largest proportion of the variance for the resistome, while the surfactants have the largest impact on the variation for the microbiota. Finally, we show that the interaction between pharmaceuticals and surfactants contributes more to the variability in the resistome than the individual compounds alone, while such interactions are less clear in the microbiota dataset (Fig. 5c and 5d).

We also collected data on the consumption of three antibiotics, ciprofloxacin, sulfamethoxazole and vancomycin, by the hospital pharmacy over the period of 2012 to 2014, that were summarized here as gram per season (Supplementary Table 3 and Supplementary Figure 3c). The studied hospital site has just been opened in the winter month of February 2012 which is probably why antibiotic consumption by the hospital pharmacy was low during winter 2012. No obvious correlation between summer peaks for the measured antibiotics in WW and their respective consumption by the hospital pharmacy was shown. (Supplementary Tables 3 and 4, and Supplementary Figure 3b and 3c). The observed peaks and variation during summer season for antibiotics, individual resistance genes and gene classes in HWW (Supplementary Figures 3a, b and 4) may be due to dry season during summer that could result in decreased flow rate of HWW.

## Discussion

In the context of globally increasing AMR, wastewaters (WW) have been identified as sources for the spread of AMR determinants (ARB, ARGs, MGEs) and chemical pollutants (often pharmaceutical residues) that may favor AMR selection during wastewater treatment and in the receiving environment. In this study we thoroughly monitored the resistome and microbiota dynamics of untreated and treated (applying conventional secondary WWT) hospital and urban WW over four years throughout the seasons in France. We identified distinct and robust resistome and microbiota signatures, in particular for untreated HWW and UWW, indicating that HWW and UWW form distinct and stable ecological niches over time. Performing machine learning (ML) classified each of the sources with high accuracy and revealed top predictive genes, gene classes and taxa for the respective WW sources before and after WW treatment (Supplementary Figure 8). Interestingly, when collapsing data-sets obtained from untreated and treated samples of the respective WW sources, ML was able to predict HWW and UWW in general with more than 93% certainty on all predictor levels (individual genes, gene classes, taxa) (Supplementary Figure 9). These top 10 predictors for HWW and UWW (Supplementary Figure 8 and 9) provide considerable marker gene classes, individual genes and taxa for the respective WW sources that indicate underlying important differences in risk to the environment and that present targets for the monitoring and management of HWW and UWW. A recent study assessed the removal efficacy of 62 Dutch wastewater treatment plants (WWTPs), applying conventional secondary WW treatment, on a selected panel of six ARGs and the gene encoding the class 1 integron integrase gene^15^. These genes (*emrB, sul1* and *sul2* (in our study synonym with *sulA*), *qnrS*, *tetM*, *blaCTX-M* and *intI*), proposed as general WW markers for risk assessment^5^, were also monitored by our resistome approach and were similarly classified as high predictors for the respective WW sources (Supplementary Figure 7, 8 and 9). All seven genes were detected in both WW sources, however in different normalized abundances (e.g. *ermB* was more abundant in UWW compared to HWW, whereas the genes *sul1, sul2, qnrS, blaCTX-M* and *intI1* were more abundant in HWW; interestingly the normalized abundance of *tetM* was comparable in both sources). We were also able to identify additional genes that could be implemented for the classification of HWW vs UWW based on their normalized abundance. For example, the streptogramin resistance gene *vatB*, and the transposase gene *ISS1N* were significantly more abundant in UWW compared to HWW, and seem to be specifically indicative for UWW.

The HWW resistome was found to be significantly enriched with resistance gene classes and genes encoding integron integrase genes compared to UWW except for the streptogramin resistance gene *vatB*. For the gene class conferring resistance to macrolides, we did not observe a significant difference between HWW and UWW; however, there is a trend for the genes encoding resistance to macrolides to have higher normalized abundance in UWW than in HWW (Figure 3a and 3b; Table 1). Based on the machine learning approach, macrolide resistance genes also contribute to the specific resistome signature for UWW (Supplementary Figure 8 and 9). Considering the fact that macrolide and streptogramin antibiotics are more frequently prescribed in the community compared to the hospital environment in France, could explain the high abundance of these gene classes in UWW^32^. In the hospital environment, antibiotics such as quinolones, beta-lactams, aminoglycosides and vancomycin are frequently used, which also could explain the relatively high abundance of gene classes conferring resistance to those antibiotics in HWW compared to UWW^32^. Furthermore, the antibiotics ciprofloxacin, sulfamethoxazole and vancomycin were detected in higher concentrations in HWW compared to UWW, which may point towards a relationship of the measured antibiotics and the detected resistome in HWW (Figure 5a and c). Interestingly, the *qnr* genes encoding quinolone resistance in HWW were the ones with the highest fold increase (161-fold) between HWW and UWW. As these genes are located on plasmids, their higher abundance in HWW indirectly reflects the likely abundance of bacteria harboring genetic elements involved in resistance dissemination as plasmids in HWW. Indeed, *qnr* genes have been described mainly in Enterobacteriales^33,34^ and we found that the relative abundance of Enterobacteriales is higher in HWW than in UWW (Figure 4).

The human gut microbiota is an important reservoir for ARGs^28,29^ and recently evidence-based data showed that the occurrence and abundance of (human) fecal pollution is a likely explanation for the detection of high amounts of ARGs in anthropogenically impacted environments^35,36^. The significant dilution of human gut bacteria in MWW observed here, is hence also likely to explain the significant reduction of the abundance of gene classes after mixing HWW with UWW.

A significant dilution of HWW-associated genes in UWW was described previously, but under circumstances that reflected a much lower rate of contribution of HWW to UWW (between 0.8 and 2.2%) at the respective study sites^9,15,19^ compared to our study site (here HWW ~33.4%). Here we show that UWW can dilute the normalized abundance of resistance genes, MGEs and integrons in untreated HWW significantly when mixed, even with an increased proportional contribution of HWW to UWW. This implies that on the scale of a small sized urbanized area as studied here, WW mixing of U and HWW bares no greater risk than separate treatment. The increase of the effluent flow rate entering in the WWTP treating the MWW due to the injected UWW may contribute largely to the observed significant dilution impact of the UWW on the HWW. Previous studies conducted on the same experimental site for shorter time periods (up to 2 years) coherently provided a detailed catalogue of pharmacological parameters, such as specific surfactants, that could further aid to discriminate UWW from HWW^27,37–39^. They also concluded that there is no greater advantage associated with separate treatment of HWW from UWW with respect to their pharmaceutical discharges and ecotoxicological impacts^27^.

With respect to the removal efficacy for ARGs and MGEs through secondary UWW treatment, we observed that the normalized abundance of six gene classes did not significantly decrease while one gene class (sulphonamides) did significantly increase (Table 2, Fig 3a and Fig 3b). The removal efficacy of secondary WW treatment in terms of absolute abundance of ARGs and ARBs is significant by reducing the overall release of ARGs and ARB into the downstream environment in general by more than 95%^15^. However, unchanged, or even increased relative or normalized abundance of ARGs and gene classes is indicative for selective processes during secondary WW treatment favoring the spread of AMR into the environment^15,40–42^. The resistome monitored here, exhibits a high mobilization potential due to the high proportion of MGEs, integrons and plasmid borne ARGs detected in all WW samples, as well as in the receiving river waters (Figure 1, MGEs and integrons account for up to 60% of the resistome of treated effluents and river waters). The MGEs, more specifically transposase genes targeted here, represent genes that are associated with the dissemination of ARGs by means of conjugative integrative elements or plasmids, across environmental bacteria and human pathogens^25,40^. There is evidence that advanced WWT such as disinfection by UV radiation or ozone treatment, or physical treatment by ultrafiltration of WW are more efficient in reducing ARB and ARGs compared to conventionally applied secondary WW treatment. However, despite reducing ARB and ARG loads significantly compared to secondary treatment often ARBs and ARGs are not completely eliminated by advanced WWT either^43^.

The exposome, a term originally coined in the context of human health epidemiology and referring to “the totality of human environmental exposures”^44^, here specifically refers to the chemical compounds quantified in our longitudinal study that represent partially the environmental or “eco-exposome”^45^ of the WW microbiota. Anthropogenic pollution through micro pollutants present in WW (particularly surfactants, antibiotics and heavy metals) has shown to have a negative impact on the environment and significantly shape the terrestrial and aquatic microbial ecosystems^8,46–49^. For example, high concentrations of antibiotics and cationic surfactants found in HWW^17,50,51^, and pharmaceutical production sites^52,53^ have been correlated with a higher abundance of ARBS, ARGs, as well as higher abundance of MGEs and integrons^17,19,54^, whereas anionic surfactants that are generally abundant in urban WW effluents (untreated and treated UWWs; “grey waters”) are associated with toxicity to aquatic and terrestrial environments^55–57^. Here, multivariate analysis (Figure 5) revealed a significant impact of the eco-exposome on the resistome and microbiota signatures of all investigated WW sources. We observed that cationic surfactants and antibiotics are specifically linked to HWW (Figure 5), which reflects the frequent use of antibiotics and active-surface agents as quaternary ammonium compounds in hospitals. Interestingly, we also found that *qac* genes that encode resistance to quaternary ammonium compounds are significantly higher in HWW than in UWW. The urban WW eco-exposome on the other hand was found to be enriched with anionic surfactants (Supplementary Table 2) which in turn were found to be specifically linked to the UWW and MWW resistome and microbiota (Figure 5). This points towards an important correlation of anionic surfactants detected in UWWs and the resistome and microbiota of UWWs, in addition to other pharmaceuticals and antibiotics. Specific measures to reduce the emission of surfactants into the environment by selective removal and improved WW treatment are currently discussed^47^ and warrant further attention. In addition, further research needs to be done to illuminate the detailed mechanism of the synergetic effects of all compounds that make up the WW eco-exposome on shaping the resistome and microbiota of WW effluents and the downstream environment.

Altogether, our findings give further emphasis to the requirement of implementing and optimizing sanitation systems and operational WWTPs on a global level, particular in countries/continents with poor water sanitation infrastructure and correlated high occurrence of multi-resistant bacteria^58^. The elimination of pollution by means of human feces, and bacterial taxa associated, through advanced or selective WW treatment, may further aid in limiting the release of ARGs associated with human gut bacteria and pathogens^35,59,60^. Finally, the data generated by this study are of important interest to policy makers concerning the risks associated with H and U WW, their putative implication into the dissemination of AMR, and provide further evidence towards the necessity of environmental pollution management in the battle of AMR and other important global health factors such as the preservation of biodiversity and the prevention of climate change^49^ (France national report: hyperlink:https://www.tresor.economie.gouv.fr/Articles/a9782706-87a4-4cdc-9a4c-86a72736315d/files/525144cc-24fb-4c93-aa5b-c67aefd3b40a).

## Methods

### Sampling and study design

126 Urban and hospital wastewater (UWW and HWW) samples were collected in Scientrier (Bellecombe WWTP), Haute-Savoie, France^27^ as part of the multi-disciplinary project SIPIBEL. The study site was implemented as an observatory for untreated and treated hospital and UWW and to evaluate their impact (during separate and subsequently mixed treatment) on the environment (e.g. the effluent receiving river). The CHAL (Centre Hospitalier Alpes Léman) hospital opened in February 2012 and includes 450 beds (140 m3/d), whereas the Bellecombe WWTP was collecting UWW of approximately 21.000 inhabitants (5200 m3/d). For more details of the SIPIBEL project, study set up, WWT and sample collection refer to Chonova et *al*. 2018 and Wiest et *al.*^27,37^, and to Figure 1. The samples included in our study were collected in monthly intervals (untreated and treated samples) by flow proportional sampling, from March 2012 through November 2015^27^. From March 2012 to December 2014, UWW and HWW were treated separately applying the same conventional (activated sludge) WWT^27,37^. Then, in the period from January 2015 through November 2015, UWW was mixed into the HWW (1:2 ratio HWW:UWW, the ratio was fixed by a local operating constraint) and added to the separate HWW treatment line resulting in a controlled mixed WW (MWW)^27^. In addition, 12 water samples of the effluent receiving river up (river upstream) and downstream (river downstream sampling point 1 and 2) of the effluent release pipes have been collected during the winter months of 2013 (January, February, November, and December). Activated sludge samples have also been collected from both sludge basins throughout the sampling campaign. Due to different resident times and flow sizes of each wastewater treatment basin, sludge dynamics for resistome and microbiota were not directly comparable hence results will not be further discussed in this study. 16S rRNA sequence data from all samples, including sludge samples, are publicly available.

### DNA isolation/ sample preparation

Water samples were filtered for microorganisms, using a filtration ramp (Sartorius, Göttingen, Germany), on sterile 47 mm diameter filter with pore size of 0.45 µm (Sartorius, Göttingen, Germany). Microorganisms were recovered from filters and subject to DNA isolation for downstream analysis, using the Power water DNA extraction kit (MoBio Laboratories Inc., Carlsbad, CA, USA). For sludge samples, 2 ml of sludge were pelleted, and DNA was extracted by following the protocol of the Fast DNA Spin kit for feces (MP Biomedicals, Illkirch, France). DNA concentration was determined by Qubit Fluoremetric Quantitation (Thermo fisher scientific, Waltham, MA USA) assays according the manufacturer’s instructions. All DNA samples were diluted or concentrated to a final concentration of 10 ng/µl for downstream qPCR and 16S rRNA analysis.

### High-throughput qPCR

Nanolitre-scale quantitative PCRs to quantify levels of genes that confer resistance to antimicrobials and heavy metals were performed as described previously^9,20^, with some modifications in the collection of primers. The primer sequences and their targets are provided in the supplementary data (Supplementary Table 5). The primer set used in the qPCR assays covered 78 genes conferring resistance to antibiotics, quaternary ammonium compounds or heavy metals. This set includes genes encoding efflux pumps (referred to as ‘efflux’, Supplementary Table 6) leading to multi-resistance at once to different antibiotic families. We also added primers for genes encoding mobile genetic elements, namely nine transposase genes^25,40^, and the class 1, 2 and 3 integron integrase genes^61^. We also included primers targeting 16SrRNA encoding DNA. Primer design and validation prior and after Biomark analysis has been done as described earlier^20^. Real-Time PCR analysis was performed using the 96.96 BioMark™ Dynamic Array for Real-Time PCR (Fluidigm Corporation, San Francisco, CA, U.S.A), according to the manufacturer’s instructions, with the exception that the annealing temperature in the PCR was lowered to 56°C. DNA was first subjected to 14 cycles of Specific Target Amplification using a 50 nM mixture of all primer sets, excluding the 16S rRNA primer sets, in combination with the PreAmp Master Mix (100-5581, Fluidigm), followed by a 5-fold dilution prior to loading samples onto the Biomark array for qPCR. Thermal cycling and real-time imaging was performed at the Plateforme Génomique GeT – INRA Transfert (https://get.genotoul.fr/en/), and Ct values were extracted using the BioMark Real-Time PCR analysis software.

### Calculation of normalized abundance and cumulative abundance

Normalized abundance of individual resistance genes was calculated relative to the abundance of the 16S rRNA gene (CTARG – CT16S rRNA) resulting in a log2-transformed estimate of gene abundance. Cumulative abundance was calculated for resistance gene classes based on the sum of the normalized abundance of individual ARGs. The differences in cumulative abundance over the indicated time periods (2012-2014; 2015 for mixed WW) are shown as an averaged fold-change ± standard deviation. The non-parametric Mann-Whitney test was used to test for significance; p values were corrected for multiple testing by the Benjamin-Hochberg procedure (Benjamini & Hochberg, 1995) with a false discovery rate of 0.05. Averaged normalized abundance data for allocated gene classes is provided in Supplementary Table 7.

### qPCR to determine absolute copy numbers of 16S rRNA genes

The qPCRs for the determination of 16S rRNA gene copy number as a proxy for the bacterial biomass was performed as described previously by Stalder et. *al*.^19^.

### 16S rRNA gene sequencing and sequence data pre-processing

Extracted DNA samples for 16S rRNA sequencing were prepared following a dual barcoded two-step PCR procedure for amplicon sequencing for Illumina. Primers of the first PCR step included universal CS1 and CS2 tags targeting the V4 region of the hypervariable region of the 16S rRNA gene using the 16SrRNA primer sequences of the earth microbiota project (http://press.igsb.anl.gov/earthmicrobiota/protocols-and-standards/16s/). During the second step of the PCR barcoded adapters suitable for multiplex illumina sequencing were added. Following pooling of the barcoded samples, the amplicon pool was cleaned to remove short undesirable fragments using Magbio HighPrep PCR beads (MagBio AC-60050), QC’ed on a High Sensitivity NGS Fragment Analyzer, and then qPCR quantified using the Kapa kit for ABI optical-cyclers. The pool was then normalized to 10nM, denatured using 0.1N NaOH followed by a 2min incubation @96C followed by 5min in an ice-water bath just prior to sequencing as per the Illumina protocol for a 2×301 MiSeq run (Illumina, Inc., San Diego, CA). DNA sequence reads from the Illumina MiSeq were demultiplexed and classified in the following manner: The Python application dbcAmplicons (https://github.com/msettles/dbcAmplicons) was used to identify and assign reads to the appropriate sample by both expected barcode and primer sequences. Barcodes were allowed to have at most 1 mismatch (hamming distance) and primers were allowed to have at most 4 mismatches (Levenshtein distance) as long as the final 4 bases of the primer matched the target sequence perfectly. Reads were then trimmed of their primer sequence and merged into a single amplicon sequence using the application FLASH^62^. Finally, the RDP Bayesian classifier was used to assign sequences to phylotypes^63^. Reads were assigned to the first RDP taxonomic level with a bootstrap score >=50.

### 16S rRNA data analysis

Illumina MiSeq forward and reverse were processed using the MASQUE pipeline (https://github.com/aghozlane/masque). Briefly, raw reads are filtered and combined followed by dereplication. Chimera removal and clustering are followed by taxonomic annotation of the resulting OTUs by comparison to the SILVA database. A BIOM file is generated that combines both OTU taxonomic assignment and the number of matching reads for each sample. Relative abundance levels form bacterial taxa (Order level) were obtained and analyzed. The obtained relative abundance OTU tables (Order level) were analyzed with Microsoft excel (Supplementary Table 8), multi-variate analysis package (see below) and by means of a machine learning approach employing a random forest algorithm (see below).

### Chemical analysis

All chemical data measured here and used for analysis where extracted from the SIPIBEL database. Solid-phase extraction (SPE) and liquid chromatography coupled with tandem mass spectrometry (LC-MS/MS) were used to measure the antibiotics ciprofloxacin, sulfamethoxazole and vancomycin and the pharmaceutical carbamazepine as detailed elsewhere^26^. Heavy metals (Zn, Cu, Ni, Pb, Cr, Gd, Hg, As and Cd) were measured with inductively coupled plasma combined with atomic emission spectroscopy (ICP-AES). Concentration of surfactants (anionic, cationic and non-ionic surfactants) were measured following standard methods approved by the French organization of standardization AFNOR as described by Wiest et. *al*.^37^.

### Multivariate analyses

Multivariate statistical techniques were used to test the influence of waste water treatment or sampling time (independent variables) on the microbiota and the resistome (dependent variables) of the different sample groups (urban, hospital, mixed), including all individual genes and genes allocated into gene classes in two independent datasets. Statistically significant influence of the treatment or the sampling time on the microbiota and the resistome were assessed by Redundancy Analysis (RDA) with 499 Monte Carlo permutations. For testing the influence of sampling time, we used sampling year or season as independent variables, and all different sample groups together (i.e., to potentially identify general patterns influencing all groups at the same time), and individually, as dependent variables. Such analysis revealed the percentage of variance that is explained by sampling time or WW treatment in each case, and whether the influence of the independent variables is statistically significant or not. The relationship between the resistome and the microbiota and the measured chemicals in the different raw water samples (eco-exposome) was visualized by means of Principal Component Analysis (PCA) biplots. Moreover, the influence of the measured chemicals (eco-exposome) on the microbiota and resistome was statistically assessed by RDA. A variation partitioning analysis was performed to assess which group of chemicals (Metals, Pharmaceuticals, Surfactants) explains the largest share of the variation of the microbiota and resistome datasets, and to explore whether the interactive effects of the groups of chemicals would have a larger influence on those datasets than the individual groups themselves. All multivariate analyses were performed with the Canoco v5.0 software^64^, using a significance level of 0.05.

### Machine learning

A Random Forest Algorithm (RFA) was used in order to predict a response variable (water sources) of each sample independently, using measurements on individual gene, gene class and microbiota level (predictor variables). To run the RFA the R-package randomForest was used: Breiman and Cutler’s Random Forests for Classification and Regression, a software package for the R-statistical environment^65^. In summary, the RFA follows the pseudo-steps: (I) the response variable and predictor variables are chosen by the user; (II) a predefined number of independent bootstrap samples are drawn from the dataset with replacement, and a classification tree is fit to each sample containing roughly 2/3 of the data, for which predictor variable selection on each node split in the tree is conducted using only a small random subset of predictor variables; (III) the complete set of trees, one for each bootstrap sample, composes the random forest (RF), from which the status (classification) of the response variable is predicted as an average (majority vote) of the predictions of all trees. Compared to single classification trees, RFA increases prediction accuracy, since the ensemble of slight different classification results adjusts for the instability of the individual trees and avoids data overfitting^66^. The Mean Decrease Accuracy (MDA), or Breiman-Cutler importance, was employed as a measure of predictor variable importance, for which classification accuracy after data permutation of a predictor variable is subtracted from the accuracy without permutation, and averaged over all trees in the RF to give an importance value [2]. It should be noted that since all predictor variables were of numeric nature, using RFA is equivalent to regression over classification trees. For the results presented here and in supplementary text, only the 2.5% of top RFA scores were considered (as presented by the resulting MDA distribution of all predictor variables), thus selecting the subset of predictor variables which appear statistically more informative than expected in the background of all predictor variables (i.e. we assume that 95% of the RFA scores fall between the 2.5th and 97.5th percentiles, as done elsewhere^67^).

### Data availability

16S rRNA sequence data are available at the European Nucleotide Archive (ENA) under the accession number PRJEB29948. All other important raw data needed to reconstruct the findings of our study are made available in the supplementary material.

## Acknowledgements

The authors thank the SIPIBEL Consortium and the SIPIBEL field observatory on the hospital’s effluents and urban wastewater treatment plants for displaying data and measurements.

## Author contributions

**C.D., M-C.P., and E.B.** designed the study, **C.D**. provided access to the SIPIBEL data collection, **M.G**. and

**E.B.** performed experiments, **E.B.**, **A.R.**, **J.L.** and **S.P.K**. performed data analysis, **L.W.** performed chemical analysis (SIPIBEL project), **E.B.**, **M-C.P.** and **C.D.** wrote the manuscript with contribution of all other co-authors.

