## Supplementary Figures 1-9 for "Classification of hospital and urban wastewater resistome and microbiota over time and their relationship to the eco-exposome" (from another version of this preprint: version 1, 2019-07-09)

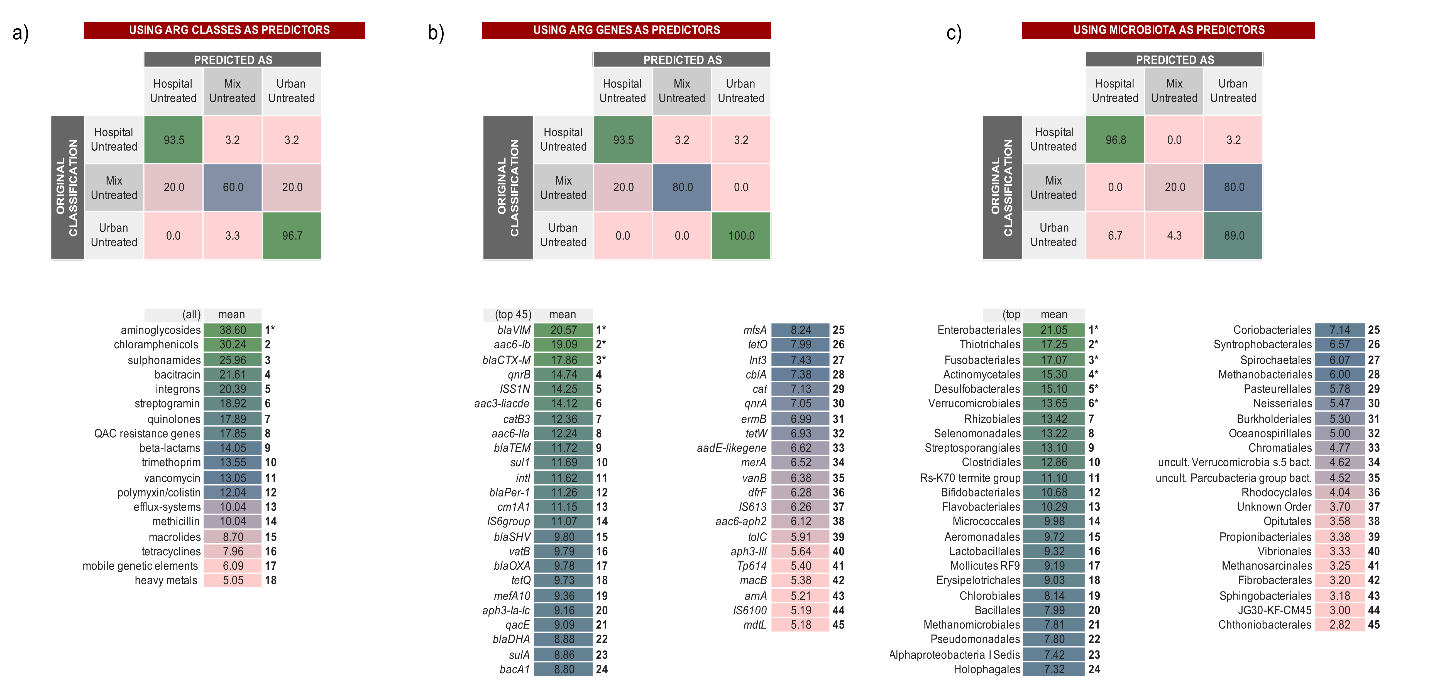

**Supplementary Figure 1. Classification success of untreated wastewater from hospital, mix and urban sources.** Top panels present the RFA classification accuracy of raw (untreated) wastewater sources when setting the predictor variables as **(a)** ARG classes, **(b)** ARG genes, and **(c)** the microbiota. The values within cells are the mean % prediction accuracy of 20 independent RFA runs with 1000 trees and default mtry. Actual wastewater source of samples in rows, predictions in columns. Colour scale is used for visualisation purposes only, from light red at 0%, blue at 50% and green at 100% of accuracy. Lower panels present the top predictor variables (capped at N=45 for visualization purposes) when setting the predictor variables as **(a)** ARG classes, **(b)** ARG genes, and **(c)** the microbiota. The values within cells are the mean MDA score of 20 independent RFA runs with 1000 trees and default mtry. Color scale is used for visualization purposes only, from light red at 0, blue at percentile 70% and green at percentile 97.25% of all MDA scores. Variables marked with * are the ones that appear more informative than expected given all the MDA scores of all predictor variables (above the cut off of percentile 97.25% of all scores, see method description).

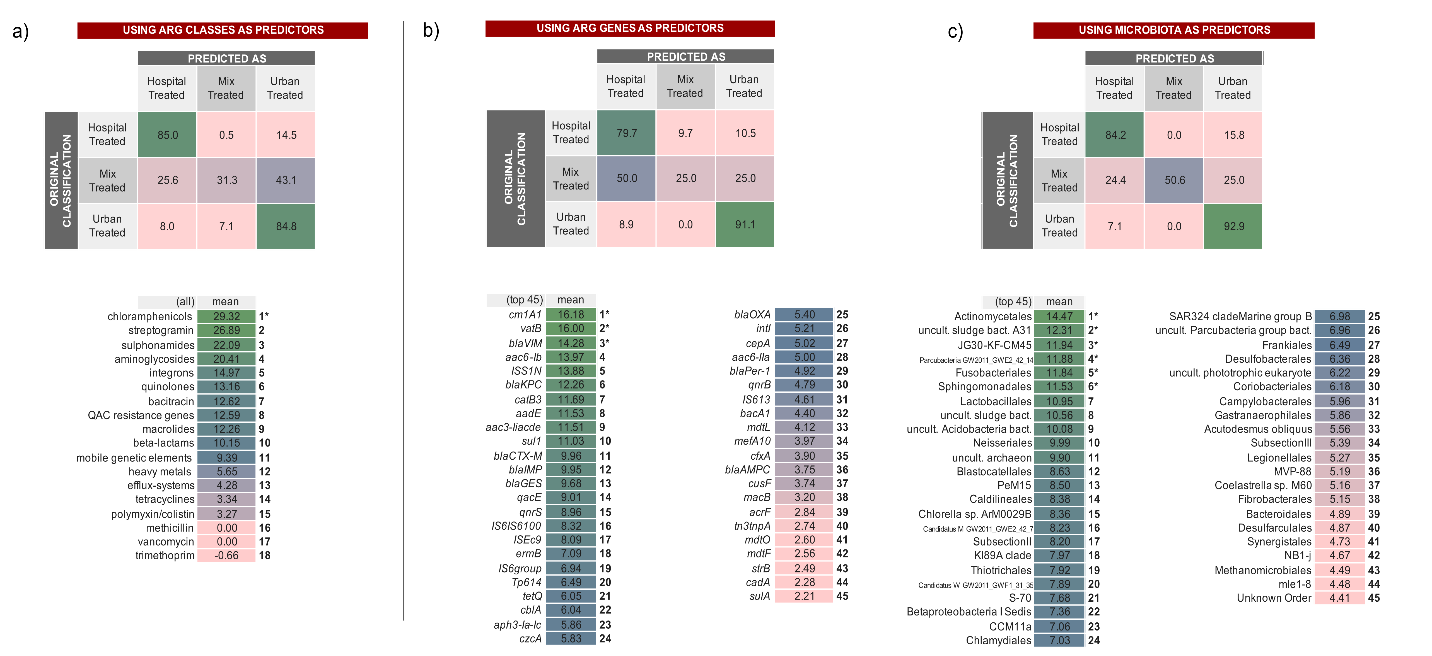

**Supplementary Figure 2. Classification success of treated wastewater from hospital, mix and urban sources.**

Upper panels present the RFA classification accuracy of treated wastewater sources when setting the predictor variables as **(c)** ARG classes, **(b)** ARG genes, and **(c)** the microbiota. The values within cells are the mean % prediction accuracy of 20 independent RFA runs with 1000 trees and default mtry. Actual water source of samples in rows, predictions in columns. Colour scale is used for visualisation purposes only, from light red at 0%, blue at 50% and green at 100% of accuracy. Lower panels present the top predictor variables (capped at N=45 for visualisation purposes) when setting the predictor variables as (a) ARG classes, (b) ARG genes, and (c) the microbiota. The values within cells are the mean MDA score of 20 independent RFA runs with 1000 trees and default mtry. Colour scale is used for visualisation purposes only, from light red at 0, blue at percentile 70% and green at percentile 97.25% of all MDA scores. Variables marked with * are the ones that appear more informative than expected given all the MDA scores of all predictor variables (above the cut off of percentile 97.25% of all scores, see method description).

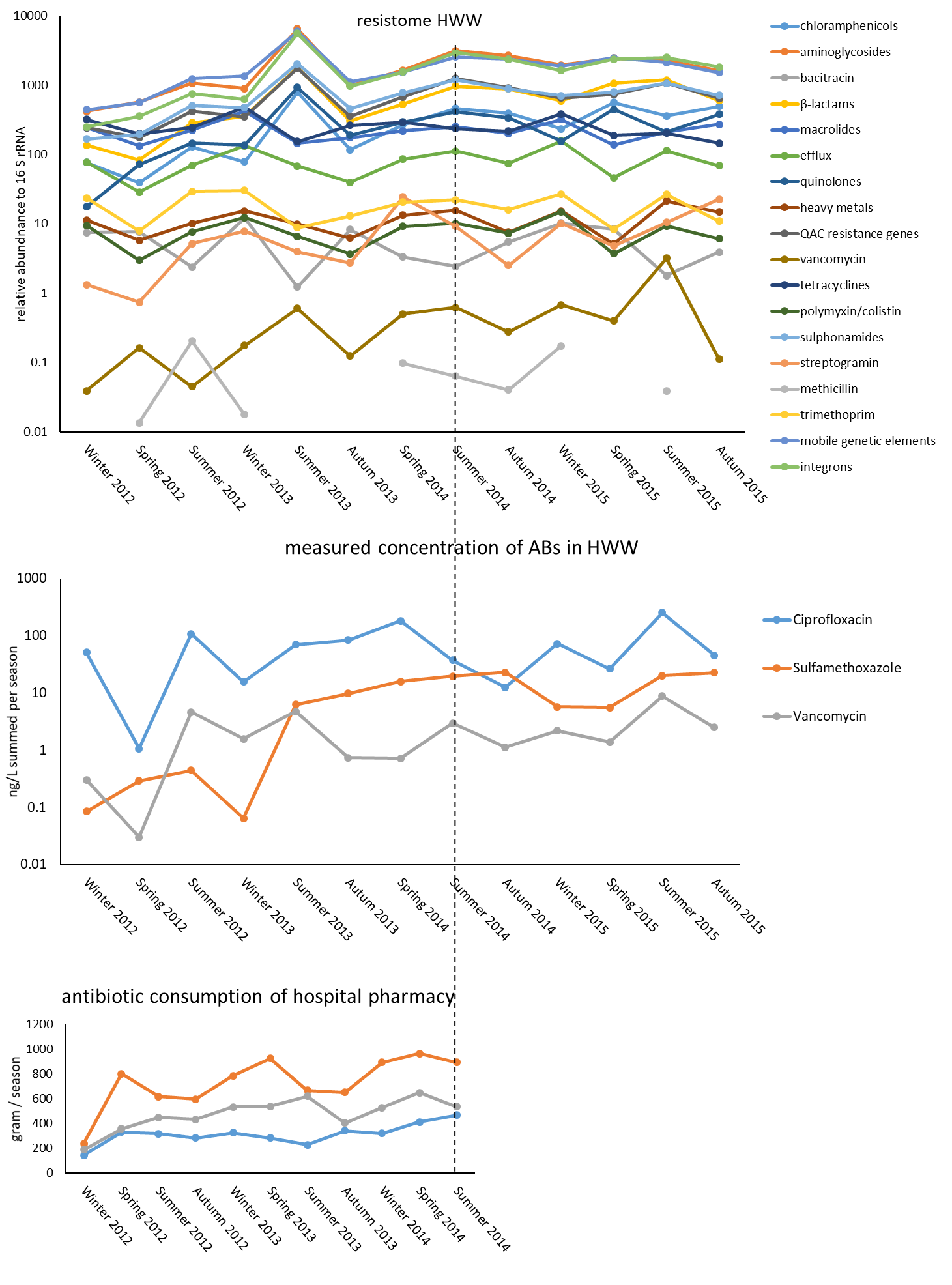
**Supplementary Figure 3**: Normalized cumulative abundance of gene classes **(a)** in relation to measured concentration of antibiotics in HWW (Ab expsosome: ciprofloxacin, sulfamethoxazole and vancomycin) **(b)** and the distribution of those antibiotics in the hospital pharmacy **(c)** per season. Vertical black line indicates summer season 2014 which represents the last data point for hospital pharmacy distribution of ciprofloxacin, sulfamethoxazole and vancomycin and facilitates direct comparisons for normalized gene class abundance and concentration of those antibiotics during the respective period in hospital WW (eco-exposome).

**Supplementary Figure 4: Redundancy analysis (RDA) exploring the relationship of the different seasons to individual genes and allocated gene classes in untreated HWW. a:** RDA for individual ARGs. Season explains 18% of the variability (MC p-value: 0.038), of which 11% is on the x-axis and 6% on the y-axis. Only the 30 genes most representative are displayed. **b**: RDA for allocated gene classes. Season explains 19% of the variability (MC p-value: 0.06, marginally significant), of which 12% is on the x-axis and 5% on the y-axis.

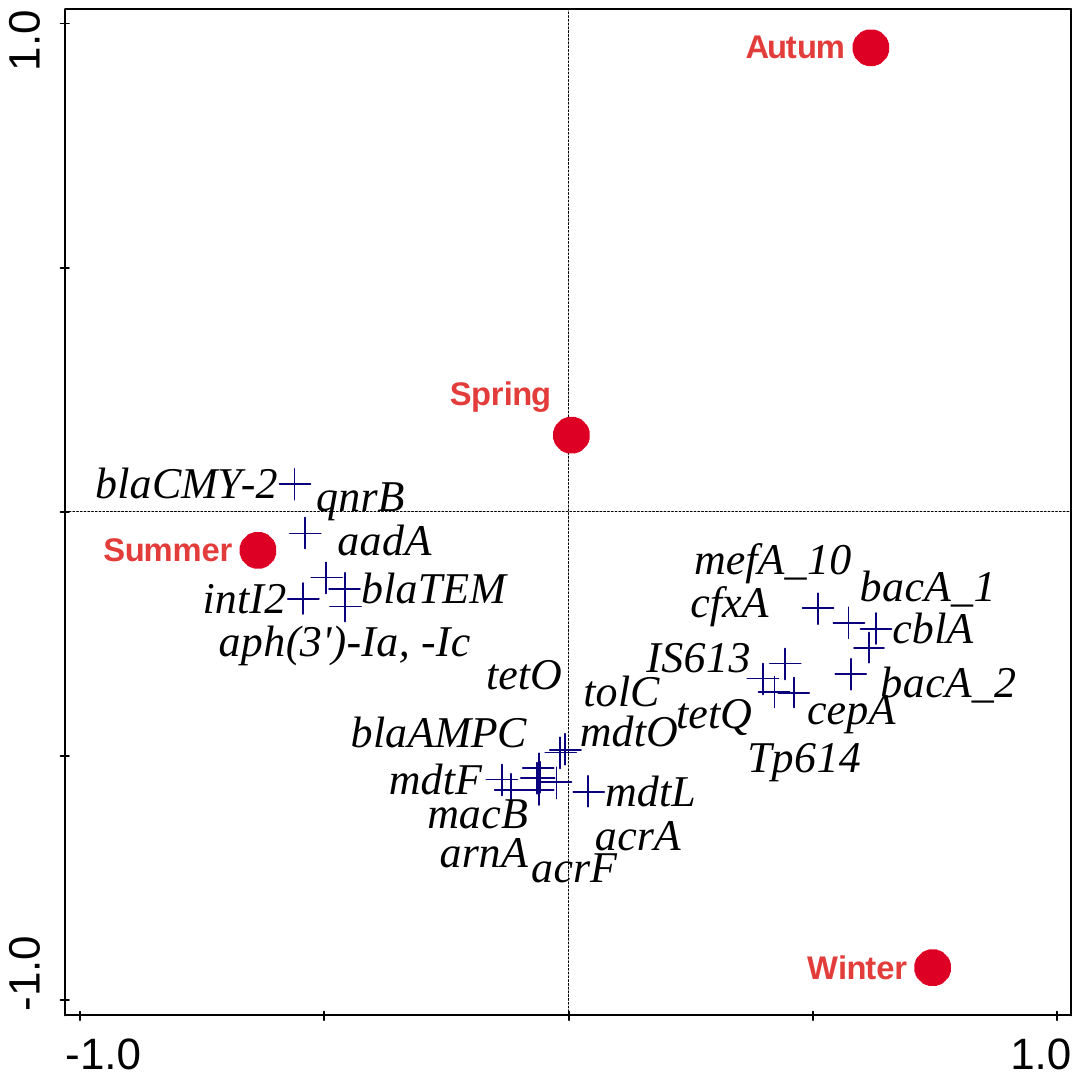

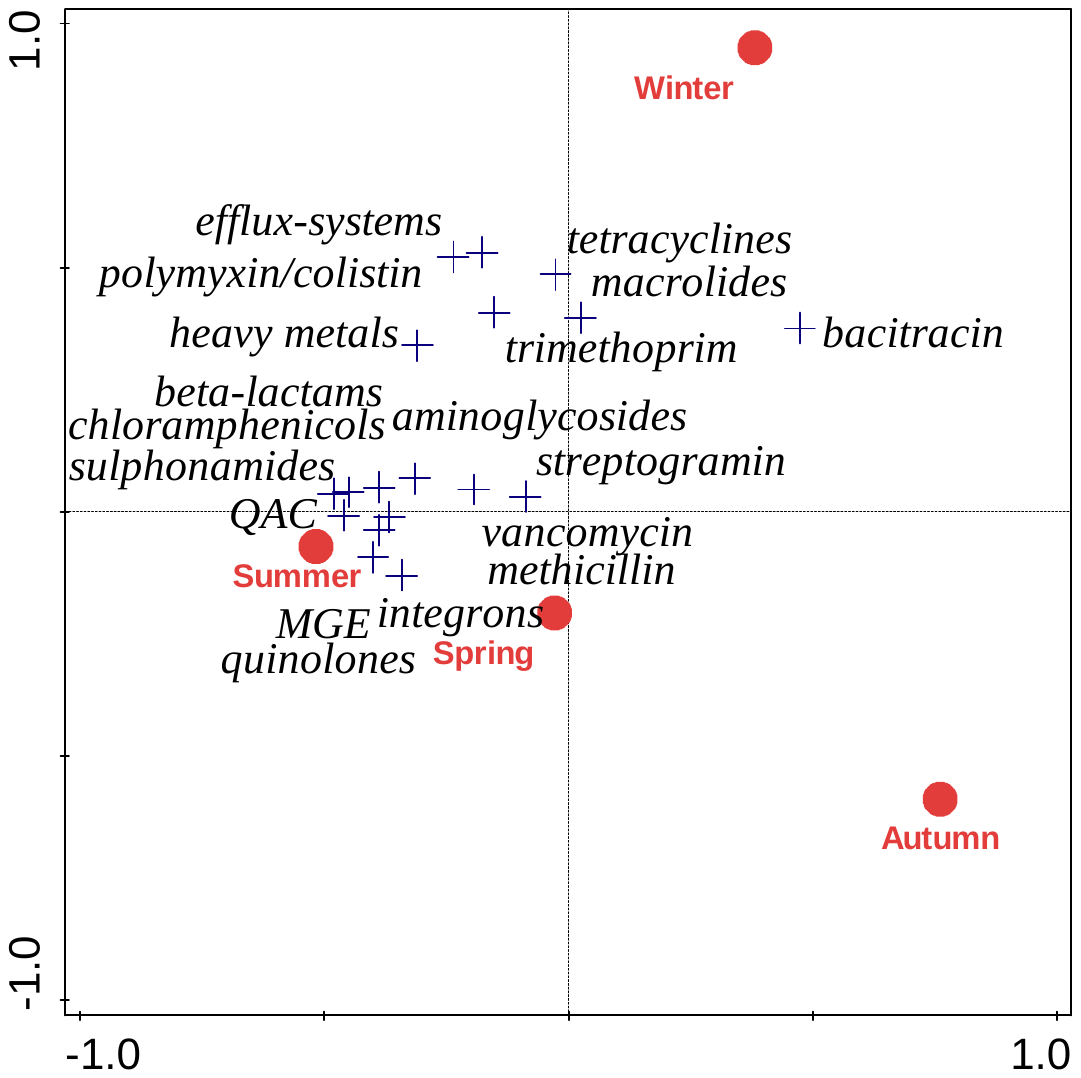

a

b

5%

6%

11%

12%

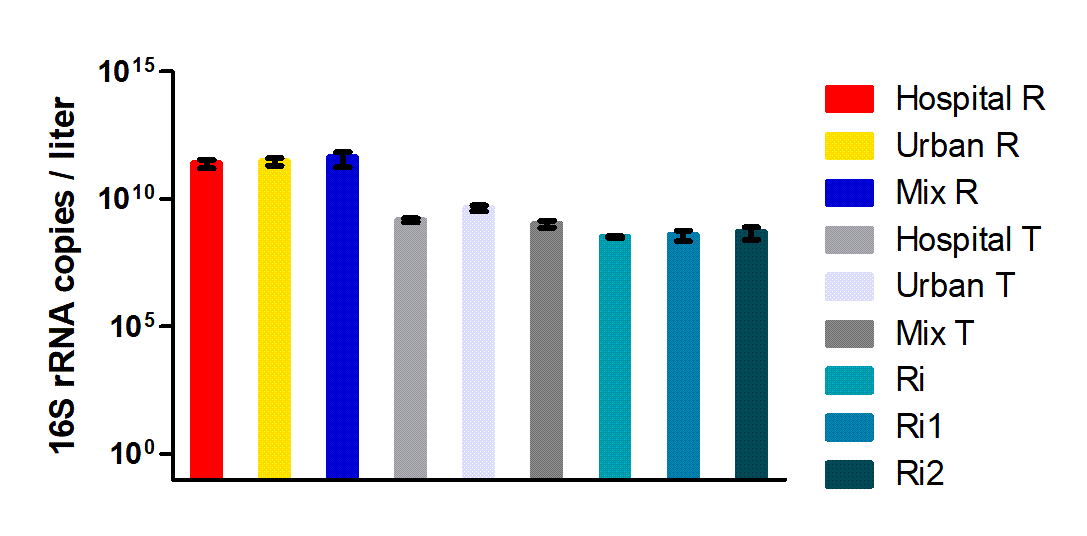

**Supplementary Figure 5:** Absolute copy number of 16S rRNA genes (Biomass; copy numbers of 16S rRNA genes / liter) for the different WW and river samples averaged over all samples) +- standard deviation. R=raw (untreated WW); T = treated WW, Ri=river upstream samples; Ri1= river downstream near effluent release pipe; River downstream of effluent release pipe.

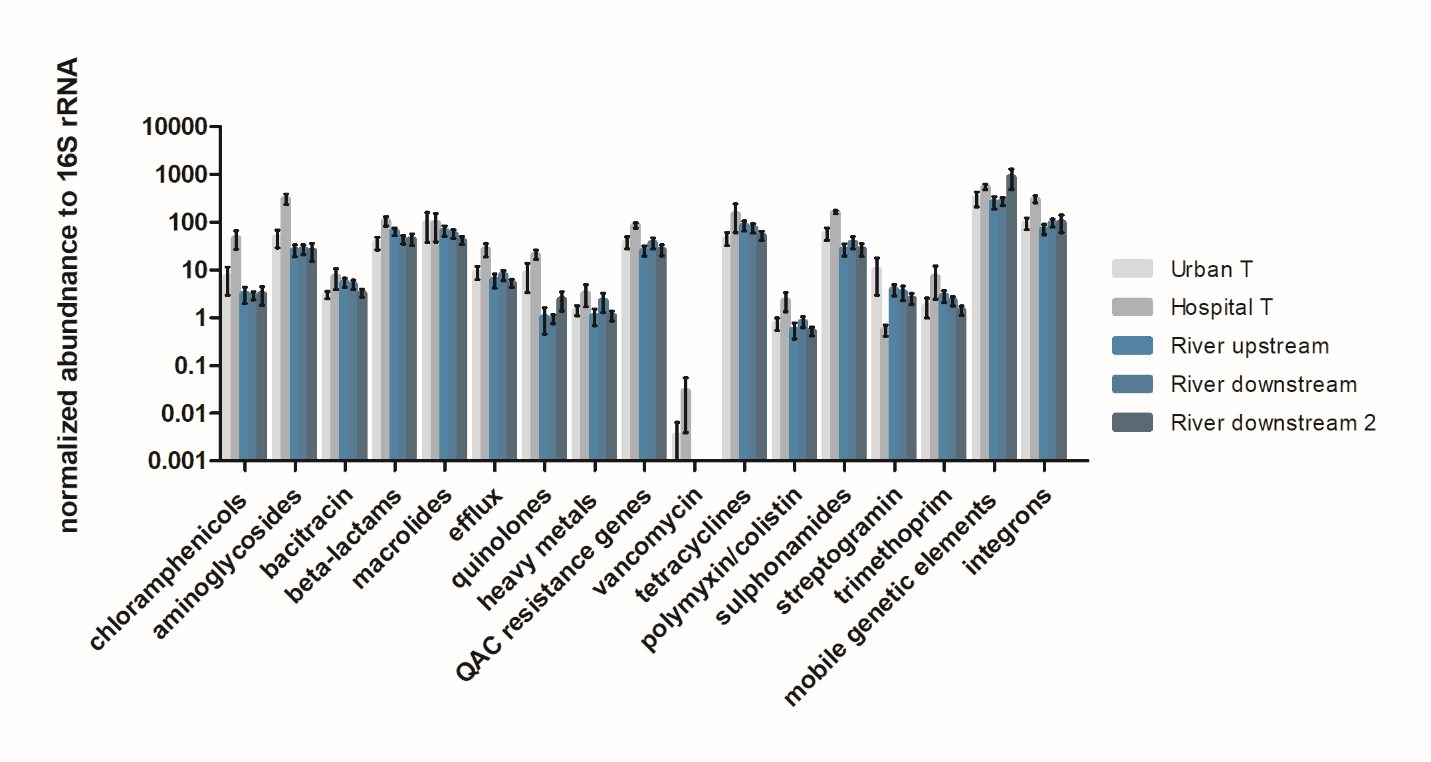

**Supplementary Figure 6: Averaged normalized abundance of ARG classes, heavy metals, MGEs and integrons over all collected river samples and their paired treated hospital and urban WW samples +- standard deviation.** Relative abundance levels of gene classes summarized over n=4 samples collected during winter season of 2013 for River and treated hospital and urban WW +- standard deviation. T= treated WW.

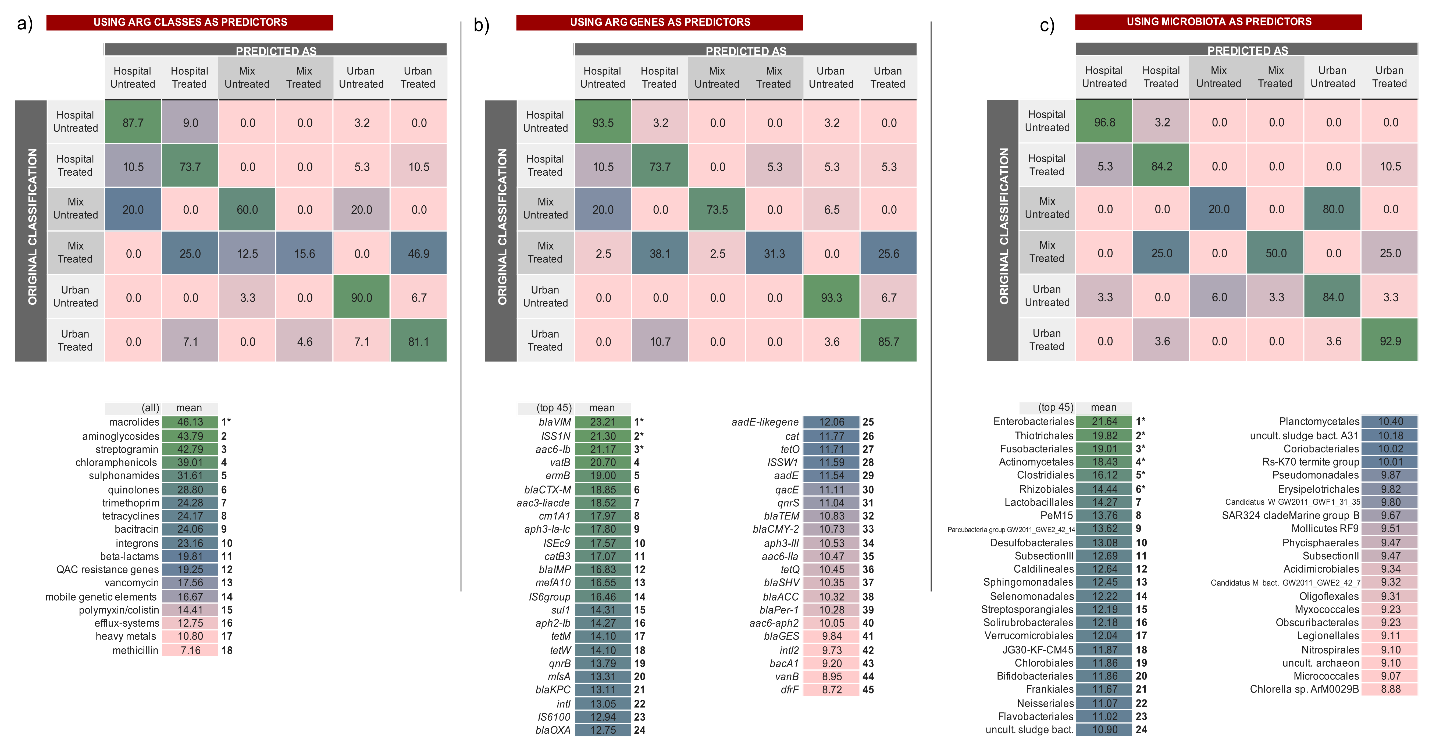

**Supplementary Figure 7. Classification success of treated and untreated wastewater from hospital, mix and urban sources**. Upper panels present the RFA classification accuracy of six water sources (Hospital Untreated / Treated, Mix Untreated / Treated, Urban Untreated / Treated) when setting the predictor variables as **(a)** ARG classes, **(b)** ARG genes, and **(c)** the microbiota. The values within cells are the mean % prediction accuracy of 20 independent RFA runs with 1000 trees and default mtry. Actual water source of samples in rows, predictions in columns. Colour scale is used for visualisation purposes only, from light red at 0%, blue at 50% and green at 100% of accuracy. Top Panels present the top predictor variables (capped at N=45 for visualisation purposes) when setting the predictor variables as **(a)** ARG classes, **(b)** ARG genes, and **(c)** the Microbiota. The values within cells are the mean MDA score of 20 independent RFA runs with 1000 trees and default mtry. Colour scale is used for visualisation purposes only, from light red at 0, blue at percentile 70% and green at percentile 97.25% of all MDA scores. Variables marked with * are the ones that appear more informative than expected given all the MDA scores of all predictor variables (above the cut off of percentile 97.25% of all scores, see method description).

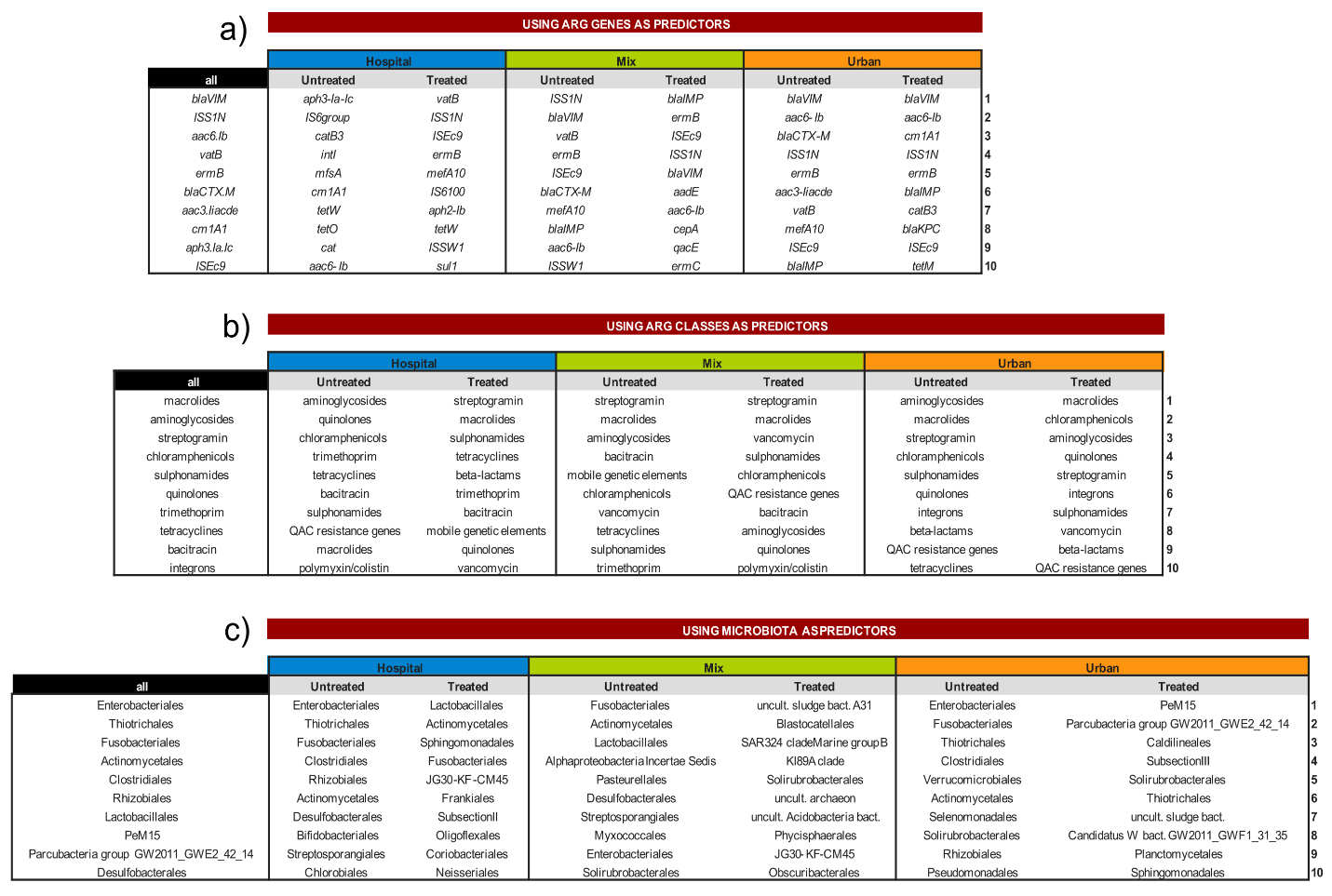

**Figure 8. Top 10 predictors for Classification success of treated and untreated wastewater from hospital, mix and urban sources.** Panels present the top 10 predictor variables for RFA classification accuracy of six water sources Hospital Untreated / Treated, Mix Raw Untreated / Treated, Urban Untreated / Treated) when setting the predictor variables as **(a)** ARG genes, **(b)** ARG classes, and **(c)** the microbiota. The column ‘all’ presents the top 10 predictors from classification exercise and accuracy presented in Supplementary Figure 7. The other columns present the top 10 predictors (from partial dependence) for particular states of the response variable (water source); Colours are solely to facilitate visualisation.

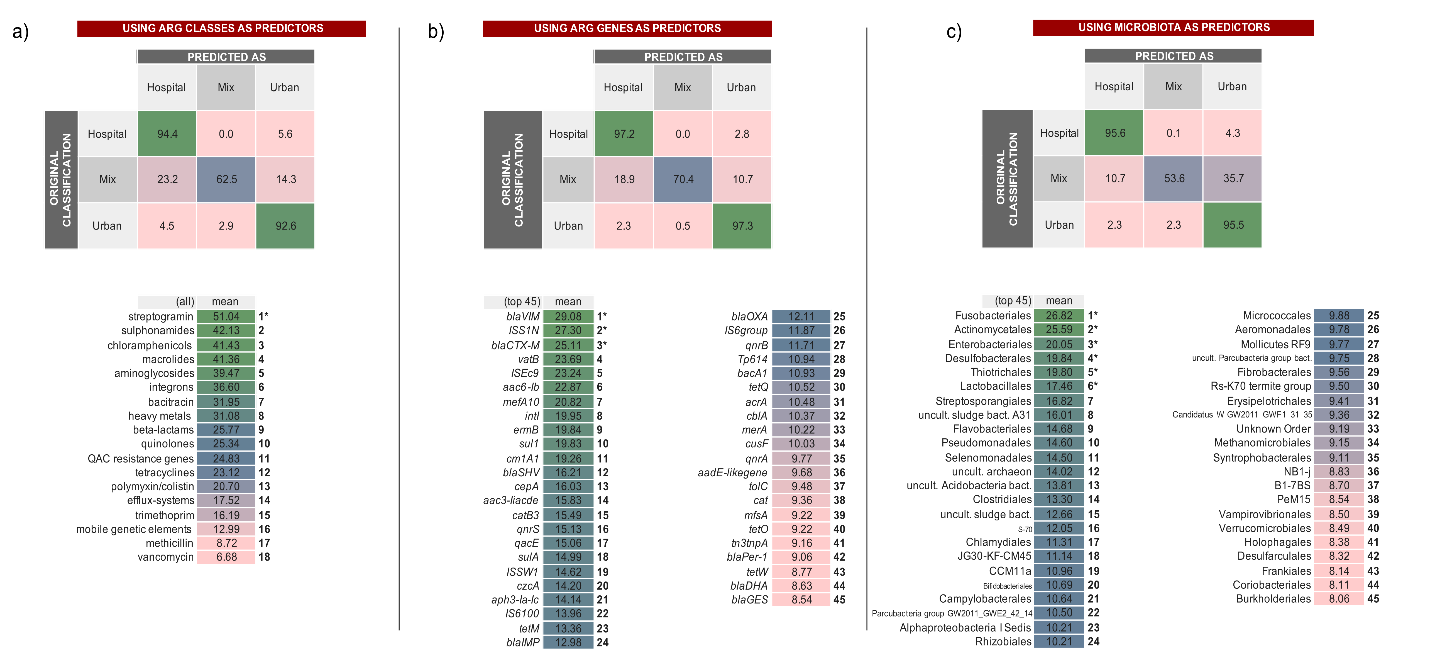

**Supplementary Figure 9.** **Classification success of hospital, mix, and urban sources when ignoring wastewater type (treated, untreated).** Upper panels present the RFA classification accuracy of three water sources (Hospital, Mix, Urban) when setting the predictor variables as **(a)** ARG classes, **(b)** ARG genes, and **(c)** the Microbiota. For this classification exercise, the original Hospital Untreated / Treated, Mix Untreated / Treated, Urban Untreated / Treated classifications of the response variable (as in Supplementary Figure 7) are aggregated to Hospital, Mix and Urban only (Untreated / Treated are collapsed). The values within cells are the mean % prediction accuracy of 20 independent RFA runs with 1000 trees and default mtry. Actual water source of samples in rows, predictions in columns. Colour scale is used for visualisation purposes only, from light red at 0%, blue at 50% and green at 100% of accuracy. Top Panels present the top predictor variables (capped at N=45 for visualisation purposes) when setting the predictor variables as (a) ARG classes, (b) ARG genes, and (c) the Microbiota. The values within cells are the mean MDA score of 20 independent RFA runs with 1000 trees and default mtry. Colour scale is used for visualisation purposes only, from light red at 0, blue at percentile 70% and green at percentile 97.25% of all MDA scores. Variables marked with * are the ones that appear more informative than expected given all the MDA scores of all predictor variables (above the cut-off of percentile 97.25% of all scores, see method description).
