## Supplementary Tables 1-4 for "Classification of hospital and urban wastewater resistome and microbiota over time and their relationship to the eco-exposome" (from another version of this preprint: version 1, 2019-07-09)

**Supplementary Table 1.** Results of the RDA analysis performed to assess the influence of sampling time (year or season) on the resistome and microbiome of the untreated (a, b) and treated (c, d) sample groups (H, U and M WW), all together and separately. Results are expressed in terms of % of explained variation in the analysis by year or season, and the calculated MC *p*-value. Significant *p-*values are in bold.

**a.** Untreated wastewater samples (sample groups analyzed together)

| MC permutations | Individual genes | | Gene classes | | Microbiome | |
| --- | --- | --- | --- | --- | --- | --- |
| H and U WW 2012-2015 | % explained | p-value | % explained | p-value | % explained | p-value |
| Year | 6.3 | 0.24 | 3.0 | 0.634 | 5.1 | 0.424 |
| Season | 4.8 | 0.42 | 11.8 | 0.874 | 5.7 | 0.282 |
| MC permutations | **Individual genes** | | **Gene classes** | | **Microbiome** | |
| U, H and M WW 2015 | % explained | p-value | % explained | p-value | % explained | p-value |
| Season | 9.4 | 0.47 | 7.9 | 0.620 | 14.7 | 0.074 |

**b.** Untreated wastewater samples (sample groups analyzed separately)

| MC permutations | Individual genes | | Gene classes | | Microbiome | |
| --- | --- | --- | --- | --- | --- | --- |
| H, U and M WW separately | % explained | p-value | % explained | p-value | % explained | p-value |
| H WW per year 2012-2015 | **23.2** | **0.004** | 12.8 | 0.242 | 11.5 | 0.26 |
| H WW per season | **18.3** | **0.038** | 18.6 | 0.058 | 15.1 | 0.072 |
| U WW per year 2012-2015 | 14.7 | 0.11 | 11.2 | 0.358 | 11 | 0.350 |
| U WW per season | 15.9 | 0.1 | 15.0 | 0.156 | 11.7 | 0.324 |
| M WW per year 2015 | Not Done | Not Done | Not Done | Not Done | Not Done | Not Done |
| M WW per season | 40.2 | 0.196 | 39.9 | 0.238 | 34.1 | 0.436 |

**c.** Treated wastewater samples (sample groups analyzed together)

| MC permutations | Individual genes | | Gene classes | | Microbiome | |
| --- | --- | --- | --- | --- | --- | --- |
| H and U WW treated 2012-2014 | % explained | p-value | % explained | p-value | % explained | p-value |
| Year | **13.9** | **0.006** | 9.0 | 0.108 | **12.9** | **0.016** |
| Season | 8.3 | 0.392 | 8.6 | 0.386 | 8.4 | 0.378 |

**d.** Treated wastewater samples (sample groups analyzed separately)

| MC permutations | Individual genes | | Gene classes | | Microbiome | |
| --- | --- | --- | --- | --- | --- | --- |
| H and U WW separately | % explained | p-value | % explained | p-value | % explained | p-value |
| H WW per year 2012-2014 | **34.9** | **0.004** | **29.5** | **0.012** | **24.2** | **0.024** |
| H WW per season | 19.5 | 0.31 | 21.3 | 0.270 | 18.5 | 0.336 |
| U WW per year 2012-2015 | **19.8** | **0.032** | 13.0 | 0.316 | **22.3** | **0.006** |
| U WW per season | 7.9 | 0.78 | 8.3 | 0.645 | 8.7 | 0.736 |
| M WW year 2015 | Not Done | Not Done | Not Done | Not Done | Not Done | Not Done |
| M WW per season | 46.3 | 0.43 | 44.9 | 0.468 | **73.7** | **0.020** |

**Supplementary Table 2:** Measured concentrations in ng/L of the eco-exposome (heavy metals, chemicals, pharmaceuticals). Values are averaged over all available samples ± standard deviation. R=raw (untreated WW); T = treated WW, Ri=river upstream samples; Ri1= river downstream near effluent release pipe; River downstream of effluent release pipe.

|  |  | Arsenic | Cadmium | Chrome | Copper | Gadolinium | Lead | Mercury | Nickel | Zinc |
| --- | --- | --- | --- | --- | --- | --- | --- | --- | --- | --- |
| Urban R | Average | 0.92 | 0.05 | 6.69 | 47.23 | 3.29 | 4.21 | 15.20 | 3.34 | 219.47 |
|  | Stdv. | 3.41 | 0.12 | 9.54 | 37.57 | 16.30 | 5.11 | 28.83 | 6.39 | 511.49 |
| Urban T | Average | 0.05 | 0.01 | 0.00 | 0.14 | 0.03 | 0.01 | 0.15 | 0.18 | 25.46 |
|  | Stdv. | 0.07 | 0.01 | 0.00 | 0.46 | 0.07 | 0.01 | 0.97 | 0.24 | 84.61 |
| Hospital R | Average | 0.46 | 0.02 | 5.69 | 67.21 | 8.56 | 1.56 | 21.00 | 1.63 | 146.55 |
|  | Stdv. | 1.87 | 0.04 | 8.49 | 33.20 | 8.84 | 2.07 | 32.99 | 4.50 | 184.24 |
| Hospital T | Average | 0.05 | 0.02 | 0.92 | 6.96 | 1.33 | 0.01 | 4.23 | 0.51 | 29.94 |
|  | Stdv. | 0.07 | 0.05 | 2.47 | 11.34 | 2.20 | 0.01 | 10.27 | 1.49 | 76.97 |
| Mix R | Average | 1.52 | 0.02 | 3.29 | 55.10 | 2.72 | 2.81 | 0.01 | 0.50 | 117.60 |
|  | Stdv. | 2.78 | 0.00 | 0.88 | 26.23 | 2.98 | 2.55 | 0.00 | 0.00 | 56.01 |
| Mix T | Average | 0.00 | 0.00 | 0.00 | 0.00 | 0.00 | 0.00 | 0.00 | 0.00 | 0.00 |
|  | Stdv. | 0.00 | 0.00 | 0.00 | 0.00 | 0.00 | 0.00 | 0.00 | 0.00 | 0.00 |
| Ri | Average | 0.89 | 0.02 | 8.43 | 1.83 | 0.05 | 0.02 | 12.68 | 2.97 | 226.56 |
|  | Stdv. | 0.91 | 0.00 | 9.10 | 2.00 | 0.00 | 0.00 | 13.72 | 4.94 | 164.22 |
| Ri1 | Average | 0.89 | 0.02 | 8.65 | 1.26 | 0.05 | 0.02 | 11.73 | 3.79 | 124.88 |
|  | Stdv. | 0.91 | 0.00 | 10.55 | 2.12 | 0.00 | 0.00 | 11.86 | 6.59 | 59.35 |
| Ri2 | Average | 1.20 | 0.02 | 7.92 | 2.90 | 0.09 | 0.02 | 10.54 | 3.31 | 165.85 |
|  | Stdv. | 1.06 | 0.00 | 8.37 | 2.70 | 0.06 | 0.00 | 9.49 | 4.87 | 66.06 |

**Supplementary Table 2:** Measured concentrations in ng/L of the eco-exposome (heavy metals, chemicals, pharmaceuticals). Values are averaged over all available samples ± standard deviation. R=raw (untreated WW); T = treated WW, Ri=river upstream samples; Ri1= river downstream near effluent release pipe; River downstream of effluent release pipe.

|  |  | Ciprofloxacin | Sulfamethoxazole | Vancomycin | Carbamazepine | Anionic surfactant | Cationic surfactant | Non-ionic surfactant |
| --- | --- | --- | --- | --- | --- | --- | --- | --- |
| Urban R | Average | 0.04 | 0.36 | 0.00 | 0.54 | 1894.67 | 145.33 | 1748.00 |
|  | Stdv. | 0.08 | 0.35 | 0.00 | 0.32 | 1179.54 | 567.24 | 2474.51 |
| Urban T | Average | 0.10 | 0.11 | 0.00 | 0.38 | 33.93 | 20.00 | 71.43 |
|  | Stdv. | 0.26 | 0.23 | 0.01 | 0.20 | 57.09 | 19.36 | 122.47 |
| Hospital R | Average | 31.98 | 4.74 | 1.07 | 1.03 | 455.56 | 551.11 | 6277.78 |
|  | Stdv. | 36.20 | 5.22 | 1.41 | 2.36 | 295.11 | 581.14 | 4326.01 |
| Hospital T | Average | 0.39 | 0.49 | 0.01 | 0.46 | 35.79 | 128.42 | 31.58 |
|  | Stdv. | 0.78 | 0.52 | 0.01 | 0.41 | 84.31 | 488.62 | 46.48 |
| Mix R | Average | 0.03 | 2.16 | 0.00 | 0.59 | 2018.00 | 532.00 | 1700.00 |
|  | Stdv. | 0.03 | 1.92 | 0.00 | 0.30 | 1081.95 | 984.00 | 715.54 |
| Mix T | Average | 1.91 | 0.87 | 0.01 | 0.64 | 50.00 | 40.00 | 100.00 |
|  | Stdv. | 1.77 | 0.38 | 0.03 | 0.29 | 0.00 | 0.00 | 0.00 |
| Ri | Average | 0.02 | 0.00 | 0.00 | 0.00 | 50.00 | 40.00 | 170.00 |
|  | Stdv. | 0.01 | 0.00 | 0.00 | 0.00 | 0.00 | 0.00 | 140.00 |
| Ri1 | Average | 0.01 | 0.00 | 0.00 | 0.01 | 50.00 | 40.00 | 372.00 |
|  | Stdv. | 0.01 | 0.00 | 0.00 | 0.00 | 0.00 | 0.00 | 389.79 |
| Ri2 | Average | 0.02 | 0.00 | 0.00 | 0.01 | 50.00 | 40.00 | 100.00 |
|  | Stdv. | 0.00 | 0.00 | 0.00 | 0.00 | 0.00 | 0.00 | 0.00 |

**Supplementary Table 3:** Estimated amount of distributed antibiotics via the hospital pharmacy during the monitoring period of 2012 to summer 2014 averaged in gram per season.

| Season | Ciprofloxacin | Sulfamethoxazole | Vancomycin |
| --- | --- | --- | --- |
| Winter 2012 | 144 | 239 | 190 |
| Spring 2012 | 330 | 801 | 358 |
| Summer 2012 | 317 | 618 | 450 |
| Autumn 2012 | 284 | 596 | 433 |
| Winter 2013 | 326 | 785 | 533 |
| Spring 2013 | 285 | 924 | 538 |
| Summer 2013 | 228 | 666 | 621 |
| Autumn 2013 | 341 | 652 | 403 |
| Winter 2014 | 321 | 892 | 528 |
| Spring 2014 | 413 | 963 | 648 |
| Summer 2014 | 468 | 893 | 535 |

| Season | Ciprofloxacin | Sulfamethoxazole | Vancomycin |
| --- | --- | --- | --- |
| Winter 2012 | 50.80 | 0.09 | 0.30 |
| Spring 2012 | 1.06 | 0.29 | 0.03 |
| Summer 2012 | 106.75 | 0.44 | 4.60 |
| Winter 2013 | 15.54 | 0.06 | 1.56 |
| Summer 2013 | 69.25 | 6.23 | 4.73 |
| Autumn 2013 | 83.33 | 9.69 | 0.74 |
| Spring 2014 | 181.15 | 15.74 | 0.71 |
| Summer 2014 | 37.33 | 19.48 | 2.94 |
| Autumn 2014 | 12.55 | 22.81 | 1.12 |
| Winter 2015 | 72.75 | 5.68 | 2.19 |
| Spring 2015 | 26.43 | 5.58 | 1.37 |
| Summer 2015 | 253.10 | 19.96 | 8.80 |
| Autumn 2015 | 44.90 | 22.41 | 2.49 |

**Supplementary Table 4:** Averaged concentration of antibiotics (ng/L) per season measured in HWW.
